# Chemotactic Migration of Bacteria in Porous Media

**DOI:** 10.1101/2020.08.10.244731

**Authors:** T. Bhattacharjee, D. B. Amchin, J. A. Ott, F. Kratz, S. S. Datta

## Abstract

Chemotactic migration of bacteria—their ability to direct multicellular motion along chemical gradients—is central to processes in agriculture, the environment, and medicine. However, studies are typically performed in homogeneous media, despite the fact that many bacteria inhabit heterogeneous porous media such as soils, sediments, and biological gels. Here, we directly visualize the migration of *Escherichia coli* populations in 3D porous media. We find that pore-scale confinement is a strong regulator of chemotactic migration. Strikingly, cells use a different primary mechanism to direct their motion in confinement than in bulk liquid. Further, confinement markedly alters the dynamics and morphology of the migrating population—features that can be described by a continuum model, but only when standard motility parameters are substantially altered from their bulk liquid values. Our work thus provides a framework to predict and control the migration of bacteria, and active matter in general, in heterogeneous environments.

**Statement of Significance:** Typical studies of bacterial motility focus on cells in homogeneous media; however, many bacteria inhabit tight porous media such as soils, sediments, and biological gels. This paper demonstrates how confinement in a porous medium fundamentally alters the chemotactic migration of *Escherichia coli*. We find that cells use a different primary mechanism to direct their motion in confinement than in bulk liquid. Further, confinement markedly alters the overall dynamics and morphology of a migrating population—features that can be described by a continuum model, but only when standard motility parameters are substantially altered from their bulk liquid values. This work thus provides a framework to predict and control the migration of bacteria, and active matter in general, in heterogeneous porous environments.

## Introduction

The ability of bacteria to migrate through tight and tortuous pore spaces critically impacts our everyday lives. For example, it can be harmful, underlying infection in the body *(1)* and food spoilage *(2)*. It can also be beneficial, enabling bacteria to deliver drugs *(3)* sense and report stimuli *(4)* protect plant roots *(5)* and degrade contaminants *(6–8)*. However, despite their potentially harmful or beneficial consequences, there is still limited understanding of how confinement in a porous medium alters the ability of bacteria to migrate: typical 3D media are opaque, precluding direct observation of cellular motion *in situ*. Thus, current understanding of migration is based on studies performed in bulk liquid.

In liquid, peritrichous bacteria swim along straight runs punctuated by rapid tumbles that reorient the cells, as established for the canonical example of *Escherichia coli (9)*. When exposed to a gradient of chemical attractant, the cells perform chemotaxis by biasing this motion. This process can mediate the directed migration of a population of cells when they continually consume a surrounding attractant: the cells collectively generate a local gradient that they in turn bias their motion along, spectacularly leading to the formation of a coherent front of cells that continually propagates *(10)*. This phenomenon can enable populations to escape from harmful environments or to colonize new terrain *(11)*. Chemotactic migration has therefore been extensively investigated under diverse conditions in bulk liquid *(10, 12–13)*.

However, confinement in a tight porous medium imposes new constraints on the ability of cells to move. For example, recent experiments have demonstrated that the paradigm of run-and-tumble motility does not describe isolated cells of *E. coli* in a gradient-free porous medium; instead, the cells exhibit a distinct mode of motility in which they are intermittently and transiently trapped between “hops” through the pore space due to interactions with the surrounding solid matrix *(14, 15)*. Hence, it is likely that confinement fundamentally alters chemotactic migration— although exactly how is unclear. Indeed, studies of microswimmers that self-propel akin to bacteria suggest that collisions with the solid matrix can suppress, or even completely abolish, coordinated motion *(16–19)*; thus, it is puzzling how coordinated multicellular migration can even occur in confined heterogeneous spaces. Nevertheless, studies in viscoelastic agar demonstrate that chemotactic migration can still arise in these heterogeneous media, although the presence of dispersed obstacles strongly hinders the ability of cells to spread over large length and time scales *(20, 21)*. Unfortunately, because such media are turbid and do not have well-defined pore structures, systematic studies of these cell-scale interactions and their impact on macroscopic migration are challenging. As a result, how confinement in a tight and tortuous space impacts chemotactic migration at the single-cell and population scales remains poorly understood. Here, we address this gap in knowledge by directly visualizing the migration of concentrated populations of *E. coli* in transparent, disordered, 3D porous media.

## Results

### Pore-scale confinement regulates, but does not abolish, chemotactic migration

We prepare porous media by confining hydrogel particles, swollen in a defined rich liquid medium with *L*-serine as the primary nutrient and attractant, at prescribed jammed packing fractions in transparent chambers. The media have three notable characteristics, as further detailed in the *Materials and Methods*. First, the packings act as rigid matrices with interparticle pores that the cells swim through (Fig. 1A, top panel), with a mean pore size *a* that can be tuned in the range ~1 to 10 μm (Fig. S1)—characteristic of many bacterial habitats. Moreover, because the hydrogel particles are highly swollen, they are freely permeable to oxygen and nutrient; thus, the influence of geometric confinement on cellular migration can be isolated and systematically investigated without additional complications arising from the influence of confinement on the spatial distribution of nutrient. Second, the media are yield-stress solids (Fig. S1); we can therefore use an injection nozzle mounted on a motorized translation stage to introduce cells into the pore space along a prescribed 3D path. As it moves through the medium, the nozzle locally rearranges the hydrogel packing and gently extrudes cells into the interstitial space; then, as the nozzle continues to move, the surrounding particles rapidly densify around the newly-introduced cells, re-forming a jammed solid matrix *(22–24)* that surrounds the population with minimal alteration to the overall pore structure *(23)* (Fig. 1A, bottom panel). This feature enables populations of bacteria to be 3D-printed in defined initial architectures within the porous media. Finally, these media are transparent, enabling tracking of fluorescent cells in 3D as they move over length scales ranging from that of single cells to that of the overall population. This platform thus overcomes three prominent limitations of common semi-solid agar assays: they do not have defined pore structures, they do not provide control over the spatial distribution of bacteria within the pore space, and their turbidity precludes high-fidelity and long-time tracking of individual cells.

**Figure 1.**
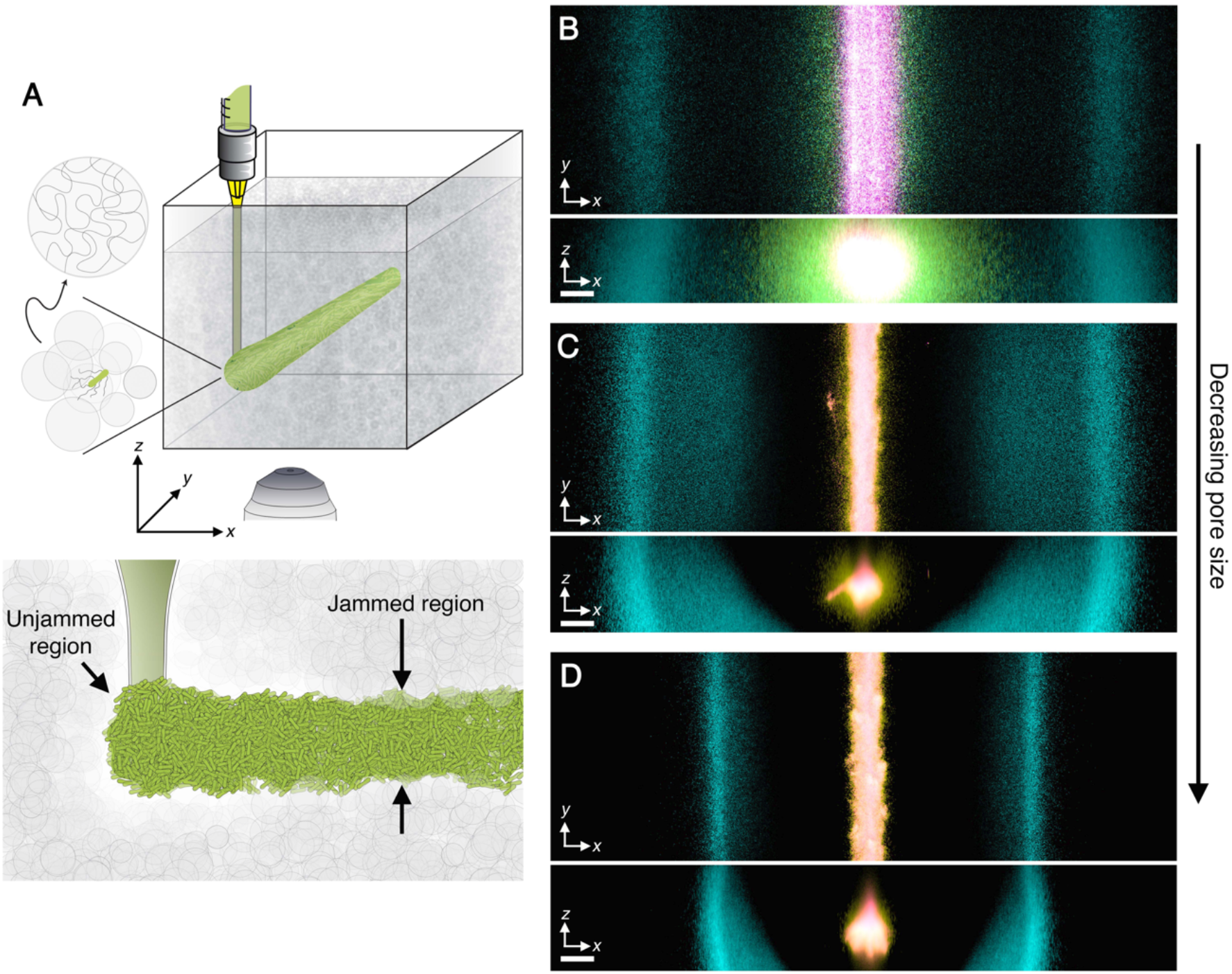
Propagating cellular fronts in porous media. **(A)** Schematic of a cylindrical population (green cylinder) 3D-printed within a porous medium made of jammed hydrogel particles (gray). The surrounding medium fluidizes as cells are injected into the pore space, and then rapidly re-jams around the cells, as shown in the lower schematic. Thus, the starting architecture of the 3D-printed population is defined by the path traced out by the injection nozzle. Each cylinder requires ~10 s to print, two orders of magnitude shorter than the duration between successive 3D confocal image stacks, ~10 min. **(B)** Top and bottom panels show bottom-up (*xy* plane) and end-on (*xz* plane) projections of cellular fluorescence intensity measured using 3D confocal image stacks. Images show a section of an initially cylindrical population at three different times (0, 1, 2.7 h shown in magenta, yellow, cyan) as it migrates radially outward in a porous medium. Panels **(C-D)** show the same experiment in media with smaller pores; (B), (C), (D) correspond to media with *a* = 2.2, 1.7, and 1.2 μm, respectively. Magenta, yellow, and cyan correspond to 0, 1.8, 10.3 h in (C) and 0, 1.3, 17.3 h in (D). All scale bars denote 200 μm; thus, a pixel corresponds to ~1 cell, indicating that the cells coherently propagate *via* multicellular fronts over length scales spanning thousands of cell body lengths.

To establish a defined initial condition, we 3D-print a ~1 cm-long cylinder of densely-packed *E. coli*, constitutively expressing green fluorescent protein throughout their cytoplasm, within a medium with *a* = 2.2 μm (Fig. 1B, magenta). The radial symmetry simplifies analysis of how the cells subsequently move. After 3D-printing, the outer periphery of the population spreads slowly (Fig. 1B, magenta-yellow and Fig. 2A, magenta-green), with a radial position *r* that varies with time *t* as ~*t*^1/2^(Fig. 2D, blue). Then, remarkably, this periphery spontaneously organizes into a ~300 μm-wide front of cells with an extended tail. This front coherently propagates radially outward (Fig. 1B, cyan; Fig. 2A, blue to cyan; Movies S1-S2), reaching a constant speed *v_fr_* ≈ 14 μm/min (Fig. 2D, blue) after an induction time *τ\** ≈ 2 h—demonstrating that coordinated multicellular migration can indeed occur in porous media. The inner region of the population, by contrast, remains fixed at its initial position and eventually loses fluorescence (Fig. S2), indicating that it is under nutrient-limited conditions.

**Figure 2.**
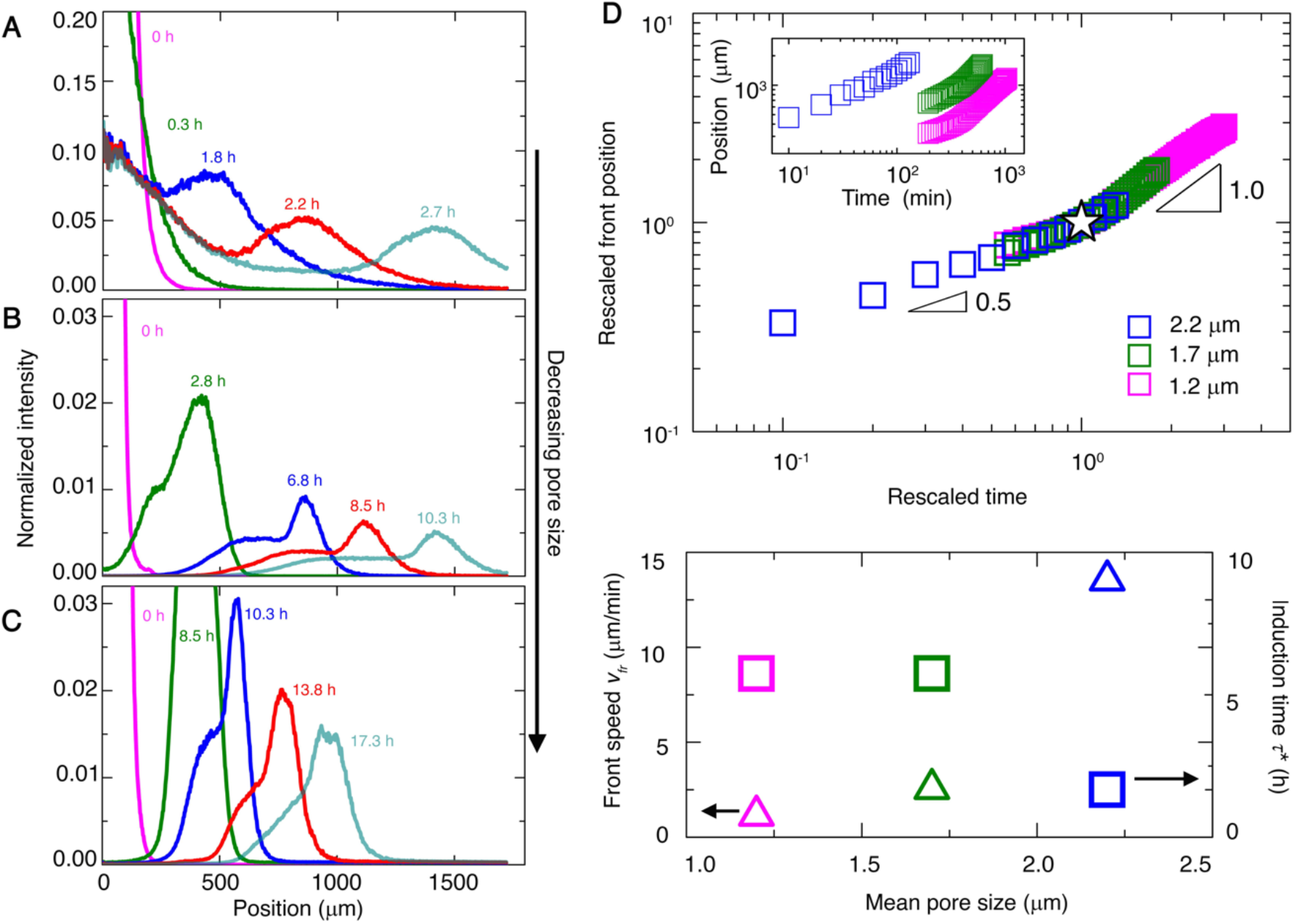
Propagation of cellular fronts is regulated by pore-scale confinement. **(A-C)** Azimuthally-averaged fluorescence intensity from cells obtained using 3D confocal stacks, normalized by its maximal initial value, for different radial positions and at different times. (A), (B), (C) show experiments performed in media with *a* = 2.2, 1.7, and 1.2 μm, respectively. In all cases, the population initially spreads outward, and then organizes into a front, indicated by the peak in the profiles, that propagates outward. **(D)** Upper panel shows leading-edge position of the propagating front over time; inset shows raw data, while main panel shows data rescaled by the crossover lengths and times (star) between diffusive and ballistic motion. Data for *a* = 1.7 and 1.2 μm begin at a later time to ensure a reliable calculation of the azimuthal average. Lower panel shows variation of front propagation speed (triangles), determined from the long-time variation of the leading-edge position, and induction time (squares), defined as the time required to transition from diffusive to ballistic motion, with mean pore size.

Without nutrient, propagating fronts do not form at all (Fig. S3). Additionally, reducing the concentration of cells in the initial population—which reduces the rate of overall nutrient consumption—increases the time required for front formation (Fig. S3). Thus, front formation is mediated by bacterial consumption of nutrient, similar to chemotactic migration in homogeneous media. However, the propagation speed is over two orders of magnitude smaller than the unconfined cellular swimming speed, and over an order of magnitude smaller than the speed of unconfined fronts *(12, 13)*: clearly pore-scale confinement regulates the dynamics of chemotactic migration.

### Individual cells bias their motion *via* a fundamentally different primary mechanism in porous media

Although the fronts of cells continually propagate outward, the individual cells do not: single-cell tracking at the leading edge of a front reveals that the cells still continue to move in all directions (Fig. 3A-B). Cells in the front exhibit hopping-and-trapping motility (Movie S3), much like isolated cells in porous media *(14, 15)*. In particular, each cell moves along a straight path of length *l_h_* within the pore space over a duration *τ_h_*—a process known as hopping—until it encounters a tight spot and becomes transiently trapped. It then constantly reorients its body until it is able to unbundle its flagella after a duration *τ_t_,* which enables it to escape and continue to hop through the pore space *(14)*. This process can be modeled as a random walk—in this case, with steps given by the hops, punctuated by pauses due to trapping (Fig. 3B).

**Figure 3.**
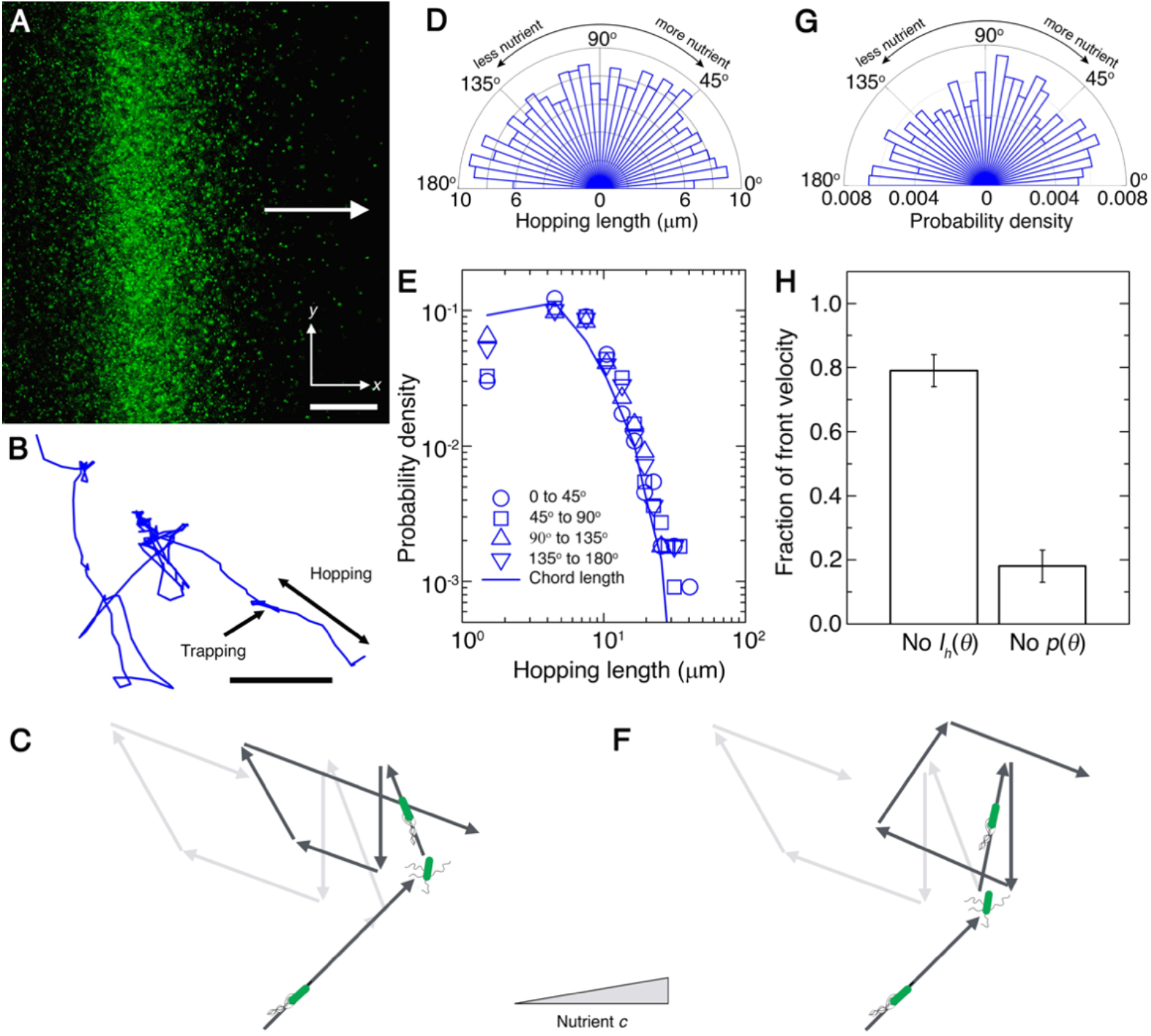
Biased motion of single cells in propagating fronts is altered by confinement. **(A)** Magnified bottom-up fluorescence intensity projection of a propagating front, showing individual cells. Arrow indicates direction of overall propagation. Scale bar denotes 200 μm. **(B)** Representative trajectory of a single cell at the leading edge of the front, over a duration of 14.9 s. The cell is localized during trapping, but moving in a directed path during hopping. Cell moves from lower right to upper left, and direction of front propagation is upward. Scale bar denotes 10 μm. **(C)** Schematic showing the primary mechanism by which cells bias their motion in bulk liquid. Light gray arrows show the unbiased random walk of a cell in gradient-free conditions; in the presence of a nutrient gradient indicated by the gray triangle, cells modulate the frequency of tumbling to bias their run length—resulting in longer runs along the direction of propagation and shorter runs in the opposite direction, as shown by the dark gray arrows. **(D)** Mean lengths of hops along different orientations |***θ***| with respect to the front propagation direction. We observe no marked directional bias: the bars are of similar length for all orientations. **(E)** Symbols show the probability density of hopping lengths along different orientations within the ranges indicated by the legend; curve shows measured chord length distribution function, which is determined by geometry, for the porous medium. The agreement between the symbols and the curve indicate that the distribution of hopping lengths is set solely by pore geometry, independent of orientation. **(F)** Schematic showing the primary mechanism by which cells bias their motion in porous media. Light gray arrows show the unbiased random walk of a cell in gradient-free conditions; in the presence of a nutrient gradient indicated by the gray triangle, cells modulate the degree of reorientation to bias their hopping orientation—resulting in more hops along the direction of propagation, as shown by the dark gray arrows. **(G)** Probability density of hopping along different orientations. We observe a directional bias: the bars are longer, indicating more hops, for orientations along the direction of front propagation, 0 ≤ |θ| ≤ 90°. **(H)** Chemotactic migration velocity calculated using Eq. 1, replacing orientation-dependent hopping lengths with the mean (first bar) or replacing orientation-dependent hopping probability with a uniform distribution (second bar). Error bars show standard deviation of velocity calculated using different angle bin widths. All data are for a = 2.2 μm.

How do these seemingly random motions collectively generate a directed, propagating front? In bulk liquid, cells detect changes in nutrient along each run, and then primarily modulate the frequency of tumbling to bias their run length—resulting in longer runs along the direction of propagation and shorter runs in the opposite direction *(9)* (Fig. 3C). However, it is unlikely that a similar mechanism could mediate migration in porous media: cells cannot elongate their hops due to obstruction by the solid matrix, nor can they shorten hops because confinement by the matrix suppresses the flagellar unbundling required to stop mid-hop *(14)*. Single-cell tracking confirms this expectation: the mean hopping lengths 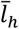 measured for hops along a given orientation *θ* relative to the direction of front propagation show no marked directional bias (Fig. 3D). The distribution of hopping lengths is instead set by pore geometry, independent of *θ*, as quantified by the chord length distribution—the probability that a straight chord of a given length *l_c_* fits inside the pore space (Fig. 3E). Hence, another mechanism must be at play.

Another mechanism also arises, albeit weakly, for chemotactic migration in bulk liquid: cells modulate the number of flagella that unbundle, and thus the degree to which their bodies reorient, during tumbling to bias the orientation of their next run *(16, 25–27)* (Fig. 3F). However, this mechanism only accounts for ~ 30% of the overall speed of front propagation in bulk liquid, with run length anisotropy accounting for ~ 70% *(13)*. Hence, why *E. coli* also employ this secondary mechanism during chemotaxis has remained a puzzle thus far.

Given that cells cannot appreciably bias their hop lengths in a porous medium, we conjecture that this putatively secondary mechanism—biasing hopping orientation—is the *primary* driver of chemotactic migration in porous media. In this mechanism, cells detect local changes in nutrient, which arise due to consumption by the entire population, along each hop. The cells then modulate their reorientation during trapping to bias the direction of their next hop along the nutrient gradient. Indeed, 90% of measured trapping events are shorter than 4 s (Fig. S4), the mean duration over which *E. coli* “remember” exposure to nutrient *(28)* suggesting that this mechanism is plausible. To directly test this hypothesis, we use our single-cell tracking to examine the probability of hopping along a given orientation, *p*(*θ*). Consistent with our expectation, we find that hops along the direction of front propagation (0 ≤ |*θ*| < 90° in Fig. 3G) are more frequent than hops in the opposite direction (90 < |*θ*| ≤ 180°). Further, we quantify the relative importance of this mechanism by computing the chemotactic migration velocity

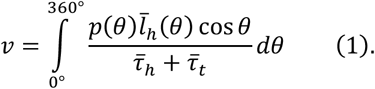

Replacing 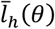 by its orientation-averaged value only changes ν by ≈ 20% (first bar in Fig. 3H)— confirming that biasing hopping length is not the primary mediator of chemotaxis, in stark contrast to the case of bulk liquid. Strikingly, however, replacing p(θ) by a uniform distribution decreases v precipitously, by over 80% (second bar in Fig. 3H)—confirming that biasing hopping orientation is the primary driver of chemotactic migration in porous media.

To further explore the influence of pore-scale confinement, we repeat our experiments in two additional media having even smaller mean pore sizes, 1.7 and 1.2 μm (Movies S4-S7). We again observe two regimes of expansion in time, with initial slow spreading followed by ballistic motion (Fig. 1C-D; Fig. 2B-C; Fig. 2D, green and magenta). Confinement is again a key regulator of these dynamics. With increasing confinement, the induction time increases while the front propagation speed considerably decreases (Fig. 2D, lower panel). The morphology of the front itself is also strongly altered by confinement: both the maximal cell density within the front, and the width of its tail, decrease with increasing confinement (Figs. 2B-C). Single-cell tracking again reveals that cells migrate by biasing hopping orientation—not by biasing hopping length, as is generally assumed (Figs. S5-S6). These effects are all missed by models of chemotactic migration in bulk liquid, in which front dynamics are determined solely by the intrinsic ability of cells to alter and respond to their chemical environment, without considering physical constraints imposed by the environment *(29)*. Other models consider environmental constraints by treating cellular motility parameters as fitting parameters or assuming their values using idealized models *(21, 30–32)*. By contrast, our experiments provide a direct way to assess how current models can be extended and applied to describe chemotactic migration in tight porous media, as detailed below.

### A continuum description of chemotactic migration requires motility parameters to be strongly altered

Our experiments reveal a clear separation of length and time scales between the biased random walks of individual cells (Fig. 3A-B) and the directed propagation of the overall front over large length and time scales (Figs. 1-2)—hinting that the macroscopic features of front propagation can be captured using a continuum description. Thus, we test whether front dynamics can be described using the classic Keller-Segel model, which is conventionally applied to chemotactic migration in bulk liquid or viscoelastic media *(11–13, 21, 29)*. Specifically, we model the evolution of the nutrient concentration *c*(*r,t*) and number density of bacteria *b*(*r,t*) *via* the coupled equations:

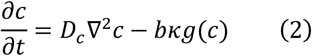

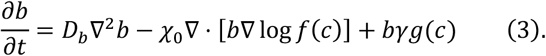

as detailed in the *Materials and Methods*. Equation 2 relates the change in c to nutrient diffusion through the medium and consumption by the population; *D_c_* is the nutrient diffusivity, *κ* is the maximal consumption rate per cell, and 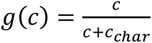 describes the influence of nutrient availability relative to the characteristic concentration *c_char_* through Michaelis-Menten kinetics. Equation 3 in turn relates the change in *b* to undirected hopping-and-trapping with diffusivity *D_b_*, biased hopping with the chemotactic coefficient *χ*_0_ and nutrient sensing function 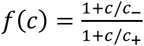, where *c*_−_ and *c*_+_quantify the range of cellular sensing, and net growth with maximal rate *γ*. Hence, this model relies on two standard quantities to describe the motion of the population over large length and time scales: the diffusivity *D_b_*, which characterizes undirected spreading, and the chemotactic coefficient *χ*_0_, which characterizes the ability of cells to bias their motion in response to a sensed nutrient gradient. In bulk liquid, their values simply depend on intrinsic cellular processes: *D_b_* is determined by the run speed and tumbling frequency *(9)* while *χ*_0_ additionally depends on properties of cellular chemoreceptors and signal transduction *(29)*. In porous media, however, confinement inhibits the ability of cells to move; it is therefore unclear whether the Keller-Segel model can describe front propagation in these more complex settings, and if so, how it must be modified.

To answer these questions, we numerically solve Eqs. 2-3 using values for all parameters estimated from direct measurements, as detailed in the *Materials and Methods—except χ*_0_, which we obtain by directly matching the asymptotic front propagation speed measured in our experiments. Importantly, we obtain *D_b_* from direct measurements of bacterial hopping lengths and trapping durations as previously established *(14)* instead of treating it as an additional free parameter or assuming its value using idealized models as is often done *(21, 30–32)*. Further, to facilitate comparison to the experiments, we determine the cellular signal—the analog of the experimentally-measured fluorescence intensity in the numerical simulations—by incorporating the fluorescence loss observed in the experiments under starvation conditions. Finally, because confinement increases the local density of cells in the pore space—increasing the propensity of neighboring cells to collide as they hop through the pore space—we explicitly account for possible cell-cell collisions that truncate both *D_b_* and *χ*_0_ at sufficiently large values of N. Because both motility parameters reflect the ability of cells to move through the pore space *via* a biased random walk with a characteristic step length *l*, we expect that they vary as ∝ *l*^2^, with lset by the mean chord length 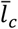 in the absence of collisions. However, when the cell density is sufficiently large, the mean distance between neighboring cells, 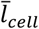 decreases below 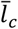; in this case, we expect that cell-cell collisions truncate *l* to ≈ 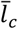. Therefore, wherever 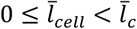, we multiply both the density-independent parameters *D_b_* and χ_0_ by the density-dependent correction factor 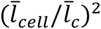.

This model indeed yields fronts of cells that form and propagate outward (solid curves in Fig. 4A-C, Movies S8-10), driven by their self-generated nutrient gradient (dashed curves); in the absence of growth, fronts still form and propagate, but their motion is hindered in a confinement-dependent manner (Fig. S8). The numerical solutions thus obtained capture the main features observed in the experiments: for all three pore sizes, the population first spreads diffusively, driven by the initially steep gradient in bacterial density, and then transitions to ballistic motion (Fig. 4D; Fig. S9) once this gradient has smoothed out (Fig. S10). Moreover, with increasing confinement, the induction time increases, while the front speed, the maximal cell density, and the width of the tail all decrease considerably (Fig. 4A-C, E)—consistent with the experimental results. As expected, the dynamics and morphologies of the fronts depend strongly on the motility parameters *D_b_* and *χ*_0_. However, unlike the case of bulk liquid, for which these parameters are set solely by intrinsic cellular processes, in tight porous media, confinement reduces these parameters by up to three orders of magnitude (Fig. 4E, bottom panel) *(11, 12)*. Further, confinement-induced cell-cell collisions play a key role in regulating chemotactic migration: in the absence of collisions, none of the simulated fronts exhibits the transition to ballistic motion observed in the experiments for any of the media tested (Fig. S9). Together, these results indicate that the Keller-Segel model can indeed describe front propagation in porous media at the continuum scale, but only when the motility parameters are substantially altered in a confinementdependent manner.

**Figure 4.**
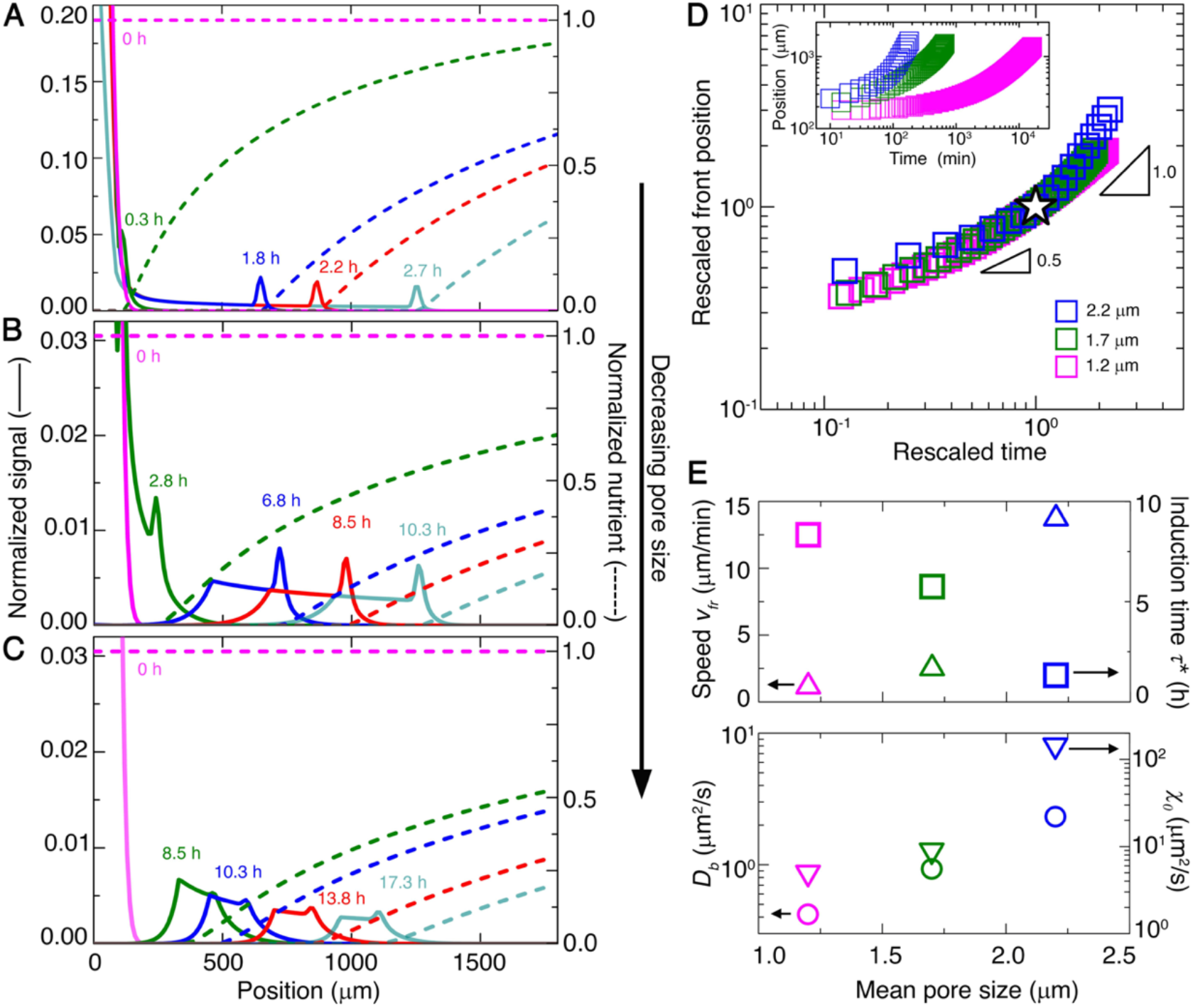
Continuum model captures dynamics of propagating cellular fronts in porous media. **(A-C)** Numerical simulations of cellular signal (solid lines) and nutrient concentration (dashed lines), normalized by maximal initial value, for different radial positions and at different times. Top to bottom panels show results for media with *a* = 2.2, 1.7, and 1.2 μm, respectively. In all cases, the population initially spreads outward, and then organizes into a front, indicated by the peak in the profiles, that propagates outward, as in the experiments. **(D)** Leading-edge position of the propagating front over time; inset shows raw data, while main panel shows data rescaled by the crossover lengths and times (star) between diffusive and ballistic motion. We observe slight deviations from the ballistic scaling for the *a* = 2.2 μm data at long times; these reflect the influence of boundaries in the system, as supported by additional simulations (Fig. S9). **(E)** Upper panel shows variation of front propagation speed (upward triangles), determined from the long-time variation of the leading-edge position, and induction time (squares), defined as the time required to transition from diffusive to ballistic motion, with mean pore size, as determined from the simulations. Lower panel shows variation of cellular diffusivity (circles), which is directly obtained from experiments, and chemotactic coefficient (downward triangles), which is determined from the simulations, with mean pore size.

### Continuum model describes long-range sensing by bacterial populations

Why can bacteria coordinate their migration in porous media, while many other microswimmers seemingly cannot? These classes of microswimmers rely on short-range interactions to coordinate their motion *(33)*. By contrast, chemotactic migration relies on the coupling between a population-generated nutrient gradient—which extends over long distances spanning hundreds of cell lengths (dashed curves, Fig. 4A-C)—and biased cellular motion along this gradient. Hence, solely through nutrient consumption, different bacteria can collectively influence and coordinate each other’s motion across long distances—even when strongly confined. For example, when the separation between two populations is smaller than the length scale ≈ 500 μm over which nutrient is depleted, they “smell” each other, and fronts only propagate away from, not toward, each other in both simulations and experiments (Fig. 5A-B, top row; Movie S11). By contrast, when the separation is much larger, fronts propagate both toward and away from each other (bottom row; Movie S12). Thus, the framework developed here provides principles to both predict and *direct* chemotactic migration.

**Figure 5.**
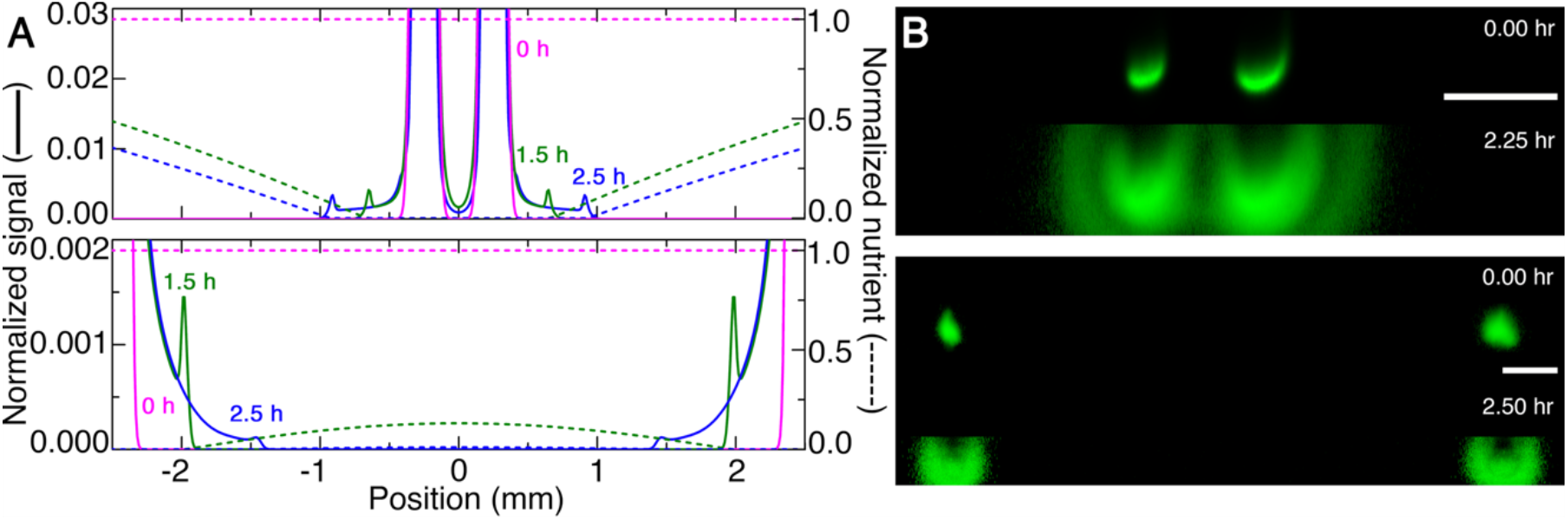
Nutrient depletion directs front propagation over long ranges. **(A)** Numerical simulations and **(B)** end-on (*xz*) fluorescence intensity projections for experiments showing front propagation from two initially cylindrical populations with axes separated by 500 μm (upper row) or 5 mm (lower row). Cells diffuse, but fronts do not propagate between the closely-separated cylinders, as shown by the cell-depleted region between the two at all times; by contrast, fronts do propagate between the further-separated cylinders. Scale bars denote 500 μm.

## Discussion

Studies of motility are typically performed in bulk liquid—even dating back to the discovery of bacteria, “all alive in a little drop of water”, in 1676. However, many bacteria inhabit tight and tortuous porous media. Our work demonstrates that chemotactic migration is fundamentally different in porous media than in bulk liquid. The paradigm of *E. coli* chemotaxis is that individual cells bias their motion primarily by modulating the frequency of reorientations, possibly with a small additional contribution due to biased reorientation amplitude. Why cells also employ this second mechanism has remained a puzzle thus far. Through direct tracking of cells performing chemotactic migration, our experiments reveal that this second mechanism is, in fact, the primary driver of chemotaxis in porous media. Thus, cells employ different mechanisms that enable them to bias their motion and forage for nutrients in different environments *(27*—motivating future studies of motility in a variety of complex settings.

Our experiments also provide a direct test of the applicability of the classic Keller-Segel model in describing chemotactic migration in highly-confining porous media. While this continuum model is broadly used for migration in bulk liquid or viscoelastic media *(11-13,21)* whether it provides a suitable description of migration in tight spaces has thus far remained unknown. Consequently, applications either utilize the Keller-Segel model *(21, 29–30, 32)* by treating both motility parameters *D_b_* and *χ*_0_ as fitting parameters, estimating them using *ad hoc* approximations, or instead turning to agent-based models that explicitly simulate the different cells, which provides tremendous insight but does not provide a continuum description *(34)*. The comparison between our experiments and simulations demonstrates that the Keller-Segel model indeed describe chemotactic migration in porous media, but only when two modifications are incorporated: (i) the cell density-independent motility parameters are reduced by several orders of magnitude from values obtained using conventional liquid assays, reflecting the hindered motion of individual cells in the tight pore space, and (ii) these motility parameters are further corrected to incorporate the influence of density-dependent cell-cell collisions, which arise more frequently in a tight pore space. Thus, pore-scale confinement is a key factor that regulates chemotactic migration, and should not be overlooked. Indeed, because the framework developed here describes migration over large length and time scales, we expect it could help more accurately describe the dynamics of bacteria in processes ranging from infections, drug delivery, agriculture, and bioremediation. Further, many other active systems—ranging from other prokaryotes, cancer cells, white blood cells, amoeba, enzymes, chemically-sensitive colloidal microswimmers, and chemical robots *(35– 41)*—also exhibit chemotaxis, frequently in heterogeneous environments and following similar rules as *E. coli*. Thus, the principles established here could be used more broadly to describe collective migration for diverse forms of active matter.

Our extension of the Keller-Segel model represents a key first step toward describing the full spatiotemporal features of chemotactic migration at the continuum scale, capturing the transition from diffusive to ballistic motion, as well as the variation of the induction time, the front speed, the maximal cell density in the front, and the width of the tail of the front with pore size observed in the experiments. However, we observe slight differences in the dynamics of the leading edge and in the shapes of the simulated fronts than those observed experimentally. These may reflect the mean-field treatment of cell-cell collisions in the model, which simplifies the details of these collisions and does not treat more sophisticated collective dynamics that arise at high local cell densities in bulk liquid *(42–46)*. Developing a more detailed treatment of these dynamics in porous media will be a useful direction for future work. Further, because our hydrogel porous media are permeable to oxygen and nutrient—similar to many biological gels, as well as many microporous clays and soils—they enable us to isolate the impact of geometric confinement on cellular migration. However, many other porous media are composed of solid matrices that are impermeable to oxygen and nutrient, resulting in more complex spatial profiles of nutrient that may further alter how cells bias their motion. Incorporating such pore-scale heterogeneities in nutrient levels will be an interesting extension of our work.

## Materials and Methods

### Preparing porous media

We prepare 3D porous media by dispersing dry granules of crosslinked acrylic acid/alkyl acrylate copolymers (Carbomer 980, Ashland) in liquid EZ Rich, a defined rich medium for *E. coli*. The components to prepare the EZ Rich are purchased from Teknova Inc. and are mixed following manufacturer directions and are autoclaved prior to use; to prepare 100 mL EZ Rich defined media, we mix 10 mL of 10X MOPS Mixture (M2101), 10 mL of 10X ACGU solution (M2103), 20 mL of 5X Supplement EZ solution (M2104), 1 mL of 20% glucose solution (G0520), and 1 mL of 0.132 M potassium phosphate diabase solution (M2102) in 58 mL of ultrapure milli-Q water. We ensure a homogeneous dispersion of swollen hydrogel granules by mixing the dispersion for at least 2 h at 1600 rpm using magnetic stirring, and adjust the pH to 7.4 by adding 10 N NaOH to ensure optimal cell viability. These granules swell further at neutral pH, resulting in a jammed medium made of ~5 to 10 μm diameter swollen hydrogel particles with ~20% polydispersity and with an individual mesh size of ~40 to 100 nm, as we established previously *(14)*, enabling small molecules (e.g. amino acids, glucose, oxygen) to freely diffuse throughout. A large volume, 4 mL, of each resulting medium is confined in a transparent-walled glass-bottom petri dish 35 mm in diameter and 10 mm in height.

### Porous medium rheology

We use shear rheology to characterize the mechanical properties of the porous media used. We load ~2 mL of each porous medium between two roughened parallel plates of 50 mm diameter separated by 1 mm in an Anton-Paar MCR301 rheometer. To determine the stiffness of the matrix, we use small-amplitude oscillatory rheology, with a strain amplitude of 1 % over a broad range of oscillatory frequencies, 0.01 −1 Hz. As shown in Fig. S1, the elastic shear moduli remain independent of frequency and are higher than the viscous shear moduli for all media tested, indicating that the porous media are elastic solids. The packings do not deform during bacterial motion, as indicated using immotile cells as tracers (Movie S3).

The porous media are yield-stress solids, as quantified by unidirectional shear measurements in which we measure the shear stress as a function of applied shear rate (Fig. S1). At low shear rates, the shear stress is constant and independent of shear rate, indicating a non-zero yield stress (~1 to 10 Pa) characteristic of a solid material. At higher shear rates, however, the shear stress follows a power law dependence on shear rate, indicating that the solid matrix becomes fluidized—the individual hydrogel particles rearrange with respect to each other and the medium yields *(23)*. This feature enables us to 3D-print populations within the pore space in defined architectures. Specifically, by applying a local stress larger than a yield stress, an injection nozzle can move through the porous medium and extrude cells into the pore space, as detailed further in the subsection “3D-printing bacterial populations” below. After extrusion of cells into the pore space, the nozzle is moved away, and the surrounding hydrogel particles rapidly re-densify around the cells, re-forming the solid matrix *(22–24)*.

### Characterizing pore space geometry

To measure the pore size distribution of each porous medium, we homogeneously disperse 200 nm-diameter carboxylated polystyrene fluorescent nanoparticles (FluoSpheres, Invitrogen) within the pore space by gentle mixing. We image the motion of the tracer particles every 34 ms using a Nikon A1R inverted laser-scanning confocal microscope with a temperature-controlled stage at 30 ± 1°C. We identify the center of each tracer using a peak finding function with subpixel precision and track the trajectory using the classic Crocker-Grier algorithm. The tracer mean-squared displacement (MSD) exhibits diffusive scaling at short length and time scales, characteristic of free diffusion within the pore space, and then transitions to subdiffusive scaling at sufficiently large length and time scales due to pore-scale confinement. Measuring the transition length scale at which the MSD becomes subdiffusive thus provides a measure of the smallest local pore dimension in the pore space; specifically, the local pore dimension *a* is given by the square root of the transition MSD plus the tracer size. Repeating this measurement for many different tracers yields the pore size distribution, which we show in Fig. S1. To measure the chord length distribution of each porous medium, we construct maximum-intensity time projections of the movies of the 200 nm tracer particle diffusion through the pore space. We then binarize these time projections into pore space and solid matrix and measure the probability that a chord—a straight line segment—fits inside the pore space. This chord length distribution is a fundamental descriptor of porous medium geometry *(14, 15)*, as further detailed in *Formulation of continuum model* below.

### 3D-printing bacterial populations

Prior to each experiment, we prepare an overnight culture of *E. coli* W3110 in LB media at 30°C. We then incubate a 1% solution of this culture in fresh LB for 3 h until the optical density reaches ~0.6, and resuspend the cells in liquid EZ Rich to a concentration of 8.6×10^10^ cells/mL. For the experiments described in the main text, we then use this suspension as the inoculum that is 3D-printed into the porous medium using an injection nozzle mounted on a motorized translation stage. As it moves through the medium, the nozzle locally rearranges the hydrogel packing and gently extrudes cells into the interstitial space; then, as the nozzle continues to move, the surrounding hydrogel particles rapidly densify around the newly-introduced cells, re-forming a jammed solid matrix *(22–24)* that compresses the cellular suspension until the cells are close-packed, similar to the case of a 3D-printed colloidal suspension *(22)*. Thus, assuming a close-packed volume fraction of 0.57 for rods of aspect ratio 4 and volume 0.6 μm^3^, we estimate the starting concentration of cells in the 3D-printed cylinders as ~0.95×10^12^ cells/mL. The 3D-printing process does not appreciably alter the properties of the hydrogel packing, as reflected in shear rheology measurements showing that different shear rates do not alter the elastic properties of the packing *(23)*. Further, this approach has been previously used to 3D-print structures made from colloidal particles of dimensions comparable to bacteria *(22)* as well structures made from mammalian cells *(24)*.

For the case of more dilute populations explored in the experiments described in Fig. S3, we dilute the suspension of cells in liquid EZ Rich to the desired starting cellular concentration by gentle mixing, which uniformly disperses the cells, into a separate sample of the jammed porous medium. We then use this medium containing cells as the inoculum that is 3D-printed into the porous medium; in this case, the 3D-printed cylinder of hydrogel particles and cells is osmotically matched to the surrounding packing of hydrogel particles and retains its shape.

To 3D-print the cells, we use a 20-gauge blunt needle as an injection nozzle, connected to a flow-controlled syringe pump that injects the inoculum at 50 μL/hr, which corresponds to a gentle shear rate of ~0.08 s^-1^ inside the injection nozzle, over two orders of magnitude lower than the shear on the cell body due to their own swimming in bulk liquid. The nozzle is mounted on a motorized translation stage that traces out a programmed linear path within the porous medium, at least ~ 500 to 1000 μm away from any boundaries, at a constant speed of 1 mm/s—resulting in a cylindrical population that provides a well-defined initial condition. Because the 3D-printed cylinders are ~1 cm long, the printing process requires ~10 s. After 3D printing, the top surface of the porous medium is sealed with a thin layer of 1-2 mL paraffin oil to minimize evaporation while allowing unimpeded oxygen diffusion. We then commence imaging within a few minutes after printing. Once a cylinder is 3D-printed, it maintains its shape until cells start to move outward through the pore space. The time needed to print each cylinder is two orders of magnitude shorter than the duration between successive 3D confocal image stacks, ~10 min. Further, because the fastest front propagates at a speed of ~14 μm/min, the overall front moves at most half a front width between each imaged time point. Therefore, the 3D printing is fast enough to be considered as instantaneous when compared with bacterial migration, and the imaging is sufficiently fast to capture the front propagation dynamics.

### Imaging bacteria within porous media

To image the motion of bacteria in 3D porous media, we use a Nikon A1R inverted laser-scanning confocal microscope at 30±1°C. To characterize front formation and propagation, we acquire a vertical stack of planar fluorescence images separated by a distance of 2.58 μm along the vertical direction, and use these to generate a 3D view of the front. We acquire these image stacks every 10-15 minutes for up to 18 h. We analyze these time-lapse image stacks using a custom MATLAB script. Specifically, we measure the azimuthally-averaged intensity of each propagating front as a function of time, only considering signal from the transverse, not the vertical (*z*), direction.

### Analysis of isolated cell motion

As we previously demonstrated *(14, 15)* isolated cells in gradient-free porous media exhibit hopping and trapping motility with hops of length *l_h_* and duration *τ_h_* punctuated by trapping events of duration *τ_t_*. To characterize this behavior, we homogeneously disperse a dilute suspension of cells (6×10^-4^ vol%) by gentle mixing within the different porous media. We visualize the cells at a fixed depth within the porous media, acquiring successive fluorescence micrographs from a slice of thickness 79 μm every 69 ms. We track 14-32 cells inside each porous medium for a minimum of 5 s. Following our previous work *(14, 15)*, we differentiate between hopping and trapping using a threshold speed of 12 μm/s, half the most probable run speed measured in bulk EZ Rich liquid: hopping is a period during which a cell moves at or faster than this threshold speed, while trapping is a period during which the instantaneous cell speed is smaller than this threshold. Although our imaging protocol yields a 2D projection of 3D cellular motion, the corresponding error in differentiating between hopping and trapping is minimal as previously established *(14)*.

### Analysis of cell motion in propagating fronts

After the 3D printed populations form propagating fronts, we image the motion of individual cells at the leading edge of the front. We chose to analyze cells at the leading edge of the front to facilitate comparison with the macroscopic measurements, which also track the leading edge of the front (as shown in Fig. 2). Indeed, studies in bulk liquid *(12)* show that the local chemotactic drift of individual cells at the leading edge matches well with the drift velocity of most of the entire front and the overall propagation speed of the entire front. Thus, chemotaxis of cells at the leading edge of the front is likely to be most representative of the overall behavior of the front. However, quantifying any systematic variation of bacterial dynamics with position in the front would be an interesting direction for future work.

We acquire successive fluorescence micrographs from a slice of thickness 79 μm every 51 ms, and track between 171 and 282 cells inside each porous medium for a minimum of 5 s. We similarly differentiate between hopping and trapping using an instantaneous speed threshold of 12 μm/s. To quantify possible directional biases, only the hopping and trapping events longer than three time points (153 ms) are included in the analysis. The angle, θ, between the direction of front propagation and the hopping direction is measured as 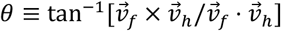 where 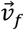 is the vector direction of front propagation and 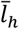 is the vector connecting the start and end point of a hop.

### Connecting single-cell motility to front propagation

Our single-cell imaging reveals that bacteria in a propagating front exhibit hopping-and-trapping motility, much like isolated cells in gradient-free porous media. Treating this process as a random walk then yields the chemotactic migration velocity *v* given by Eq. 1 of the main text; *θ* represents the hopping angle with respect to the direction of macroscopic front propagation and thus *l_h_* cos *θ* represents the projected length of a hop.

Specifically, each hop identified using imaging of single cells at the leading edge of the front yields a measurement of *θ, l_h_*(*θ*), and *τ_h_*, while each trapping event yields a measurement of *τ_t_*. We directly calculate *p*(*θ*) using all measurements of *θ*, and we calculate 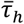 and 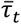 by averaging over all hopping and trapping measurements. We calculate 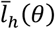 by averaging the measured *l_h_* over all hops having *θ* within a bin spanning (*θ − δθ,θ + δθ*). Then, we calculate *v* as a discrete sum: 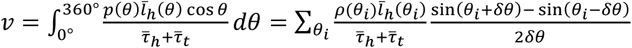 where the sum is over all bins 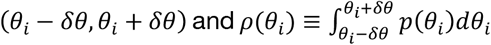 represents the fraction of all hops having orientations within a specified bin. To ensure our choice of bin width 2*δθ* has no effect on the results, we vary the bin width from 45 degrees to the smallest value for which each bin contains at least 20 data points, corresponding to 4, 2.61, and 10 degrees for the media with mean pore size 2.2, 1.7, and 1.2 μm, respectively. The velocity results for different bin widths are shown in Fig. S5. The calculated velocity overshoots the actual front velocity due to limitations in tracking very long trap times, thus artificially lowering the average trap time in the discrete sum and raising the velocity. However, these plots demonstrate that the order of the three conditions tested—uniform *p*(*θ*) uniform 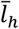, or both *p(θ)* and *l_h_* being *θ*-dependent—is consistent across all bin widths. The variation in velocity for different bin widths is reported in the standard deviation shown in the bar charts of main text Fig. 3. Replacing *p(θ)* by a uniform distribution decreases *v* precipitously, confirming that biasing hopping orientation—presumably by modulating the number of flagella that unbundle during trapping, and thus the amplitude of cell body reorientation, as has been analyzed previously *(13, 25–27)*—is the primary driver of chemotactic migration in porous media. The results thus obtained are not sensitive to the presence of hops spanning the boundary of the field of view; removing hops beginning in a buffer region 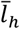 wide on all boundaries, yet keeping hops that end in this region, still yields similar results to those presented here. Further, we note that a similar analysis performed for cells at varying positions throughout the front will be an interesting direction for future work; our analysis only focuses on cells at the leading edge of the front to facilitate comparison with the macroscopic tracking of the leading edge shown in Fig. 2. Further, as suggested by others for experiments in bulk media *(12)* the local chemotactic drift of individual cells near the leading edge matches well with the drift velocity of most of the entire front and indeed, the overall speed of the entire front.

### Formulation of continuum model

To mathematically model front propagation, we build on previous work *(11–13, 29, 47–53)* to describe the evolution of the nutrient concentration *c*(*r, t*) and number density of bacteria *b*(*r,t*) *via* Eqs. 2-3. The continuum model, which is conventionally applied to chemotactic migration in bulk liquid or viscoelastic media, relies on two standard quantities to describe the motion of the population over large length and time scales: the diffusivity, which characterizes undirected spreading, and the chemotactic coefficient, which characterizes the ability of cells to bias their motion in response to a sensed nutrient gradient. Our single-cell tracking in the absence of a nutrient gradient provides a direct determination of the diffusivity, which we then use directly as an input to the model. Our single-cell tracking at the leading edge of the chemotactic front also demonstrates the importance of cellular reorientation bias in driving chemotaxis; however, the single-cell data do not yield a direct determination of the chemotactic coefficient, because this quantity also depends on properties of cellular chemoreceptors and signal transduction, as well as the exact nutrient conditions, all of which are unknown. Therefore, as is conventionally done, we determine this parameter by directly fitting the long-time speed predicted by the continuum model to the experimentally-determined front speed. The continuum model does not explicitly incorporate the exact mechanism by which cells bias their motion; it simply requires a bias in cellular motion, as confirmed by the single-cell tracking.

#### Choice of *c*(*r, t*)

The medium contains 10 mM *L*-serine as the most abundant nutrient source and attractant *(54)*. Moreover, *E. coli* consume *L*-serine first in mixed media *(55)* and are known to respond most strongly to serine as a chemoattractant compared to other components of the media we use *(56–60)* as well as compared to oxygen *(59)*. Thus, unlike other work exploring mixtures of different nutrients and attractants *(11)* in our experiments *L*-serine acts as the primary nutrient source and attractant. When the primary nutrient and primary attractant are different chemical species, metabolically active cells continue to grow and divide in the wake of the propagating front *(11)*—unlike in our experiments, for which the inner region of the population remains fixed and eventually loses fluorescence, indicating that it is under nutrient-limited conditions *(60)*. We therefore focus on *L*-serine in the continuum model, represented by the concentration field *c*(*r, t*).

We note that while *L*-serine can exhibit toxicity at high concentrations *(61)* consumption by the cells reduces the local nutrient levels by over one to two orders of magnitude within the propagating fronts themselves (indicated in Fig. 4A-C of the main text); thus, we do not expect or see any indication of possible toxicity of *L*-serine in the experiments.

Our numerical simulations focus on the nutrient concentration *c*(*r, t*); however, incorporating oxygen concentration as an additional field variable, initially at 250 μM throughout *(62)* that diffuses *(63)* with diffusivity 2500 μm^2^/s and is consumed by the bacteria at a maximal rate of 1.2 ×10^-12^ mM(cell/mL)^-1^s^-1^ and with a characteristic Michaelis-Menten level *(12)* of 1 μM reveals that the oxygen profile is remarkably similar to that of the nutrient (Movie S13): oxygen becomes depleted in the same region as the nutrient, consistent with the idea that the front contains aerobically metabolically active cells while behind the front cells are deprived of both nutrient and oxygen.

We note that the nutrient levels of our liquid medium are nearly two orders of magnitude larger than the levels under which *E. coli* excrete appreciable amounts of their own chemoattractant *(64)*. Moreover, chemoattractant excretion results in the collapse of cells into pointlike aggregates *(64-66)* which are not observed in our experiments. Thus, under the nutrient-rich conditions explored in our work, it is unlikely that bacteria in the front excrete appreciable levels of their own chemoattractant.

For all of these reasons, our model incorporates a single nutrient and attractant through the field *c*(*r, t*) for simplicity.

#### Nutrient diffusion

Molecules of *L*-serine (size ~1 nm) are nearly two orders of magnitude smaller than the hydrogel particle mesh size ~ 40-100 nm. Moreover, the *L*-serine isoelectric point is 5.7, lower than our pH of 7.4, and the polymers making up the hydrogel are negatively charged under our experimental conditions; we therefore do not expect that attractive electrostatic interactions or complexation arise. Thus, we do not expect that steric or electrostatic interactions with the hydrogel matrix impede *L*-serine diffusion, and we take the nutrient diffusivity *D_c_* to be equal to its previously measured value in pure liquid, 800 μm^2^/s.

#### Nutrient consumption

The total rate of nutrient consumption is given by *bκg*(*c*) where *κ* is the maximum consumption rate per cell and *g*(*c*) = *c*/(*c+c_char_*) describes the influence of nutrient availability through Michaelis-Menten kinetics i.e. it quantifies the reduction in consumption rate when nutrient is sparse as established previously *(11, 21, 67–69)* with *c_char_* = 1 μM as determined previously *(70)*. We use a value of *κ* = 1.6 × 10^−11^mM(cell/mL)^-1^s^-1^ comparable to values determined previously *(21)* that yields values of *v_fr_* that match the experimental values and yields front peak heights that match experimental values of the cellular signal.

#### Cellular diffusivity

Porous confinement alters both the undirected and directed components of cell motion. Our previous work *(14)* showed that isolated cells move in an undirected manner *via* hopping-and-trapping motility, with a cellular diffusivity *D_b_* that characterizes motion over time scales much larger than *τ_t_* ~ 1-10 s. To determine *D_b_* for each porous medium, we use experimental measurements of the hopping lengths *l_h_* and trapping durations *τ_t_* of isolated cells in gradient-free conditions in each porous medium, and calculate 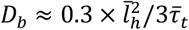, where the factor of 0.3 is an empirical correction determined previously *(14)*. We finally obtain *D_b_* = 2.32, 0.93, and 0.42 μm^2^/s for porous media with a = 2.2, 1.7, and 1.2 μm, respectively.

#### Cellular chemotaxis

We employ an advective term −Δ · (*bv_c_*) to describe biased motion along a chemoattractant gradient, where the chemotactic velocity 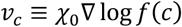 quantifies the abilities of individual cells to logarithmically respond to the local nutrient gradient. Specifically, the function 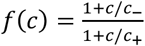 established previously *(11)* quantifies the ability of the cells to sense nutrient levels *(71–77)* where *c*_−_ = 1 μM and *c*_+_ = 30 μM are the upper and lower bounds of logarithmic sensing, and the chemotactic coefficient *χ*_0_ quantifies the ability of bacteria to bias motion in response to the sensed nutrient gradient. Although heterogeneity in *χ*_0_ may be present within the population *(12)* we focus our analysis on the effect of pore size by assuming all individual bacteria have identical chemotactic capabilities. Since our experiments demonstrate that the ability to bias motion is dependent on pore-scale confinement, we use *χ*_0_ as the pore size-dependent fitting parameter. We vary *χ*_0_ to match the numerically-simulated long-time front speed with that of the experiment. We finally obtain *χ*_0_ = 145, 9, and 5 μm^2^/s for porous media with *a* = 2.2, 1.7, and 1.2 μ m, respectively.

#### Influence of cellular crowding

Both motility parameters *D_b_* and *χ*_0_ reflect the ability of cells to move through the pore space *via* a biased random walk with a characteristic step length *l*. In unconfined liquid, 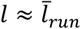, the mean run length. For the case of isolated cells in gradient-free porous media, 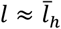, the mean hopping length. Hops are runs that are truncated by obstruction by the solid matrix, and thus the hopping lengths *l_b_* are determined solely by the geometry of the medium—specifically, by the lengths of straight paths that fit in the pore space as demonstrated previously *(14)* with 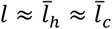, the mean length of a straight chord that fits in the pore space *(14, 15, 78)*. However, our work here focuses on highly concentrated bacterial populations in which cell-cell collisions can become appreciable—specifically, when the mean distance between cells, 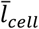, becomes smaller than 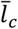. In this case, cell-cell collisions further truncate *l*, and instead 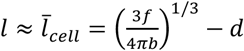, where *f* is the volume fraction of the pore space between hydrogel particles, *b* is the local bacterial number density, and *d* = 1 μm is the characteristic size of a cell. Thus, wherever *b* is so large that 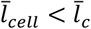, we multiply both *D_b_* and *χ*_0_ in Eqs. 2-3 by the correction factor 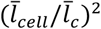 that accounts for the truncated *l* due to cell-cell collisions. Moreover, wherever *b* is even so large that this correction factor is less than zero—i.e. cells are jammed—we set both *D_b_* and *χ*_0_ to be zero. Based on our experimental characterization of pore space structure *(15)* we use *f* = 0.36, 0.17, and 0.04, and 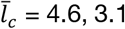, 3.1, and 2.4 μm, for porous media with *a* = 2.2, 1.7, and 1.2 μm, respectively.

#### Cell growth

To obtain the cell doubling time *τ_2_*, we measure the first division time for isolated cells within a gradient-free small-pore medium that inhibits cellular motion. Specifically, we measure the duration between the first cell division and the second cell division for 13 cells to find the average cell division time to be *τ_2_*= 60 minutes. The rate at which cells grow is then given by *bγg*(*c*) where *γ = ln(2)/τ_2_* is the maximal doubling rate per cell and *g*(*c*) again describes describes the influence of nutrient availability through Michaelis-Menten kinetics i.e. it quantifies the reduction in growth rate when nutrient is sparse. Because *c* and *b* are coupled in our model, we do not require an additional “carrying capacity” of the population to be included, as is often done *(11, 21)*: we track nutrient deprivation directly through the radially-symmetric nutrient field *c(r, t)*.

#### Loss of cellular signal

We experimentally observe that while the periphery of a 3D-printed population forms a propagating front, the inner region remains fixed and eventually loses fluorescence, indicating that it is under nutrient-limited conditions. Specifically, the fluorescence intensity of this fixed inner population remains constant for *τ_detay_* = 2 h, and then exponentially decreases with a decay time scale τ_starve_ = 29.7 min (Fig. S2). We incorporate this feature in our numerical simulations to determine the cellular signal, the analog of measured fluorescence intensity in the numerical simulations. Specifically, wherever *c*(*r′,t′*) drops below a threshold value, for times *t > t′ + τ_delay_*, we multiply the cellular density *b*(*r′,t*) by 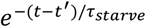, where t′ is the time at which the position *r′* became nutrient-depleted. This calculation yields the cellular signal plotted in Figs. 4-5 of the main text, for which delayed fluorescence loss yields the zigguratlike shape of the propagating front.

### Implementation of numerical simulations

To numerically solve the continuum model, we use an Adams-Bashforth-Moulton predictor corrector method where the order of the predictor and corrector are 3 and 2, respectively. Since the predictor corrector method requires past time points to inform future steps, the starting time points must be found with another method; we choose the Shanks starter of order 6 as described previously *(79, 80)*. For the first and second derivatives in space, we use finite difference equations with central difference forms. Time steps of the simulations are 0.01 s and spatial resolution is 10 μm. Because the experimental chambers are 3.5 cm in diameter, we use a radial distance of 1.75 × 10^4^μm for the size of the entire simulated system.

To match the symmetry of a single 3D printed cylinder, we use a one-dimensional axisymmetric coordinate system with variation in the radial coordinate. To simulate two 3D printed lines (Fig. 5 of the main text), we instead use a one-dimensional Cartesian coordinate system (“slab” geometry) that avoids the unnecessary use of two spatial coordinates while still demonstrating the key features of the experiment. No flux boundary conditions are used for symmetry in the center and at the walls of the simulated region for both field variables *b* and *c*.

The initial cylindrical distribution of cells 3D-printed in the experiments has a diameter of 100 ± 10 μm; so, in the numerical simulations, we use a Gaussian with a 100 μm full width at half maximum for the initial bacteria distribution *b*(*r,t* = 0), with a peak value that matches the 3D-printed cell density in the experiments, 0.95 × 10^12^ cells/mL. The initial condition of nutrient is 10 mM everywhere, characteristic of the liquid media used in the experiments. The initial nutrient concentration is likely lower within the population initially due to nutrient consumption during the 3D printing process; however, we find negligible effects of this initial condition on the characteristics of front propagation (Movie S14).

To assess convergence of the numerical solutions, we perform simulations with varying spatial and temporal resolution. Even for the case of the largest pore size medium, which has the largest value of *χ*_0_/*D_b_* and thus requires the finest resolution, we find the long-time front speed obtained with spatial resolution of 10 μm is within ~14% that obtained with a resolution of 5 μm—in close agreement—and the bacterial profiles *b*(*r,t*) have similar characteristics. For the intermediate pore size medium, we find that the long-time front speed obtained with spatial resolution of 10 μm and temporal resolution of 0.01 s is within ~ 5% the value obtained with spatial resolution of 5 μm and temporal resolution of 0.001 s, and the bacterial profiles *b*(*r,t*) have similar characteristics (Fig. S13), confirming that the resolution is sufficiently fine so that our results are not sensitive to the choice of resolution.

### Comparison between simulations and experiments

The goal of our modeling is to identify the essential physics needed to extend the classic Keller-Segel model to the case of heterogeneous porous media, with minimal alteration to the input parameters. We therefore do not expect perfect quantitative agreement between the experiments and simulations. Instead, we hope that our work will motivate future extensions of the model that provide an even better match to the experiments, as further detailed below.

Overall, we find good agreement between the simulations and experiments. Specifically, in all cases we observe a comparable crossover from diffusive to ballistic motion, with comparable induction times, front speeds, and front peak heights, indicating that our simplified extension of the Keller-Segel model provides an essential first step toward capturing the dynamics of chemotactic migration in porous media at the continuum scale. However, we do observe discrepancies between the model and the experiments. These discrepancies likely reflect (i) the influence of boundaries in the experiments, (ii) the simplified treatment of cell-cell collisions, (iii) differences in the values of the exact parameters input to the simulations, as detailed further below.

#### (i) The influence of boundaries in the experiments

While the experiments initially have cylindrical symmetry, with the initial 3D-printed cylinder placed far from all boundaries, as fronts propagate, they begin to approach the bottom boundary of the imaging chamber. Specifically, the simulations indicate that the region of nutrient depletion reaches the bottom boundary after ~0.5-1 h for experiments in the largest pore size media; in this case, the symmetry of the fronts is no longer cylindrical in the experiments, but has a rectilinear component. We conjecture that this feature gives rise to the deviation in the long-time scaling in the experiments performed in the largest pore size media, indicated by the blue circles in the figure on the previous page. To test this conjecture, we have repeated the simulations, but in rectilinear coordinates; the leading-edge position of the propagating front over time for simulations performed in rectilinear coordinates is shown in the top panel of Fig. S9; colors correspond to those in Fig. 4 of the main text. Stars indicate the crossover from diffusive to ballistic motion. In this case, we observe closer agreement to the ballistic scaling observed in the experiments than simulations performed in cylindrical coordinates, confirming our conjecture.

#### (ii) The simplified treatment of cell-cell collisions

Because confinement increases the local density of cells in the pore space—increasing the propensity of neighboring cells to collide as they hop through the pore space—in the model, we explicitly account for possible cell-cell collisions that truncate both motility parameters at sufficiently large values of the cell density. We do this using a mean-field treatment, in which both motility parameters are truncated by a density-dependent geometric correction factor. A key finding of our work is that incorporating cell-cell collisions is essential in the model; neglecting them entirely, as is conventionally done, yields fronts that do not achieve the measured ballistic scaling for any of the porous media tested (as shown in the bottom panel of Fig. S9). However, this mean-field treatment simplifies the details of these collisions, assuming that they simply truncate hops and do not alter trapping, and also does not treat more sophisticated collective dynamics that arise at high local cell densities in bulk liquid, such as swarming. Developing a more detailed treatment of these dynamics in porous media will be a useful direction for future work.

#### (iii) Differences in the values of the exact parameters input to the simulations

While our simulations use values for all input parameters estimated from our and others’ direct measurements, the values used may not exactly match those corresponding to the experiments, given the uncertainty inherent in determining these parameters (e.g., the maximal nutrient consumption rate, the characteristic nutrient level in the Michaelis-Menten function). Thus, the simulations may not perfectly reproduce the experiments. For simplicity, we fix the values of these parameters using previous measurements and focus instead on the variation of the motility parameters with pore size. We anticipate that our findings will help to motivate future work that better constrains the values of the input parameters to the Keller-Segel model.

## Acknowledgments

It is a pleasure to acknowledge Tommy Angelini for providing microgel polymers; Average Phan and Bob Austin for providing fluorescent *E. coli;* the Stone lab for use of the Anton-Paar rheometer; and Bob Austin, Zemer Gitai, Yannis Kevrekidis, Jasmine Nirody, Josh Shaevitz, Howard Stone, Sankaran Sundaresan, Ned Wingreen, and Ricard Alert for stimulating discussions. This work was supported by NSF grant CBET-1941716, the Project X Innovation fund, a distinguished postdoctoral fellowship from the Andlinger Center for Energy and the Environment at Princeton University to T.B., support from the Keller Center REACH program for F.K., the Eric and Wendy Schmidt Transformative Technology Fund at Princeton, and in part by funding from the Princeton Center for Complex Materials, a Materials Research Science and Engineering Center supported by NSF grant DMR-1420541. This material is also based upon work supported by the National Science Foundation Graduate Research Fellowship Program (to J.A.O.) under Grant No. DGE-1656466. Any opinions, findings, and conclusions or recommendations expressed in this material are those of the authors and do not necessarily reflect the views of the National Science Foundation.

## Author Contributions

T.B. and S.S.D. designed the experiments; T.B. performed all experiments; D.B.A., J.A.O., F.K., and S.S.D. designed the numerical simulations; D.B.A. performed all numerical simulations; T.B., D.B.A., and S.S.D. analyzed the data; S.S.D. designed and supervised the overall project. All authors discussed the results and implications and wrote the manuscript.

## Supplementary Information

### Supplementary Movie Captions

**Movie S1 (separate file): Bottom-up view of chemotactic migration in media with largest pores.** Maximum intensity projection of chemotactic migration from a cylinder of close-packed *E. coli* 3D-printed within a medium with *a* = 2.2 μm.

**Movie S2 (separate file): End-on view of chemotactic migration in media with largest pores.** Maximum intensity projection of chemotactic migration from a cylinder of close-packed *E. coli* 3D-printed within a medium with *a* = 2.2 μm.

**Movie S3 (separate file): Single cell hopping and trapping at the leading edge of a propagating front.** Bottom-up imaging of cells at the leading edge of a front propagating in a medium with *a* = 2.2 μm; direction of front propagation is upward.

**Movie S4 (separate file): Bottom-up view of chemotactic migration in media with medium sized pores**. Maximum intensity projection of chemotactic migration from a cylinder of close-packed *E. coli* 3D-printed within a medium with *a* = 1.7 μm.

**Movie S5 (separate file): End-on view of chemotactic migration in media with medium sized pores.** Maximum intensity projection of chemotactic migration from a cylinder of close-packed *E. coli* 3D-printed within a medium with *a* = 1.7 μm.

**Movie S6 (separate file): Bottom-up view of chemotactic migration in media with smallest pores.** Maximum intensity projection of chemotactic migration from a cylinder of close-packed *E. coli* 3D-printed within a medium with *a* = 1.2 μm.

**Movie S7 (separate file): End-on view of chemotactic migration in media with smallest pores.** Maximum intensity projection of chemotactic migration from a cylinder of close-packed *E. coli* 3D-printed within a medium with *a* = 1.2 μm.

**Movie S8 (separate file): Numerical simulation of chemotactic migration in media with largest pores.** Numerical simulations of cellular signal (blue lines) and nutrient concentration (red lines), normalized by maximal initial value, for different radial positions and at different times, for a medium with *a* = 2.2 μm.

**Movie S9 (separate file): Numerical simulation of chemotactic migration in media with medium sized pores.** Numerical simulations of cellular signal (blue lines) and nutrient concentration (red lines), normalized by maximal initial value, for different radial positions and at different times, for a medium with *a* = 1.7 μm.

**Movie S10 (separate file): Numerical simulation of chemotactic migration in media with smallest pores.** Numerical simulations of cellular signal (blue lines) and nutrient concentration (red lines), normalized by maximal initial value, for different radial positions and at different times, for a medium with *a* = 1.2 μm.

**Movie S11 (separate file): End-on view of chemotactic migration from two closely-spaced cylindrical populations.** Maximum intensity projection of chemotactic migration from two cylinders of close-packed *E. coli* 3D-printed within a medium with *a* = 1.7 μm and with axes separated by 0.5 mm. Chemotactic fronts only move outward, not inward between the cylinders.

**Movie S12 (separate file): End-on view of chemotactic migration from two well-spaced cylindrical populations.** Maximum intensity projection of chemotactic migration from two cylinders of close-packed *E. coli* 3D-printed within a medium with *a* = 1.7 μm and with axes separated by 5 mm. Chemotactic fronts move both outward and inward between the cylinders.

**Movie S13 (separate file): Numerical simulation of oxygen levels during chemotactic migration in media with medium size pores.** Numerical simulations of cellular signal (blue lines) and oxygen concentration (red lines), normalized by maximal initial value, for different radial positions and at different times, for a medium with *a* = 1.7 μm. The resulting oxygen profile is qualitatively similar to the nutrient profile shown in Movie S9. Both oxygen and nutrient are depleted behind the front, supporting our expectation that the front contains aerobically metabolically active cells while behind the front cells are starved.

**Movie S14 (separate file): Numerical simulation of chemotactic migration in media with medium size pores, with initial nutrient depletion.** Numerical simulations of cellular signal (blue lines) and nutrient concentration (red lines), normalized by maximal initial value, for different radial positions and at different times, for a medium with *a* = 1.7 μm. In this case, the nutrient is not saturated everywhere initially, but is instead depleted within the initial population; however, we find identical results to the case with nutrient saturated everywhere.

### Supplementary Figures

**Figure S1.**
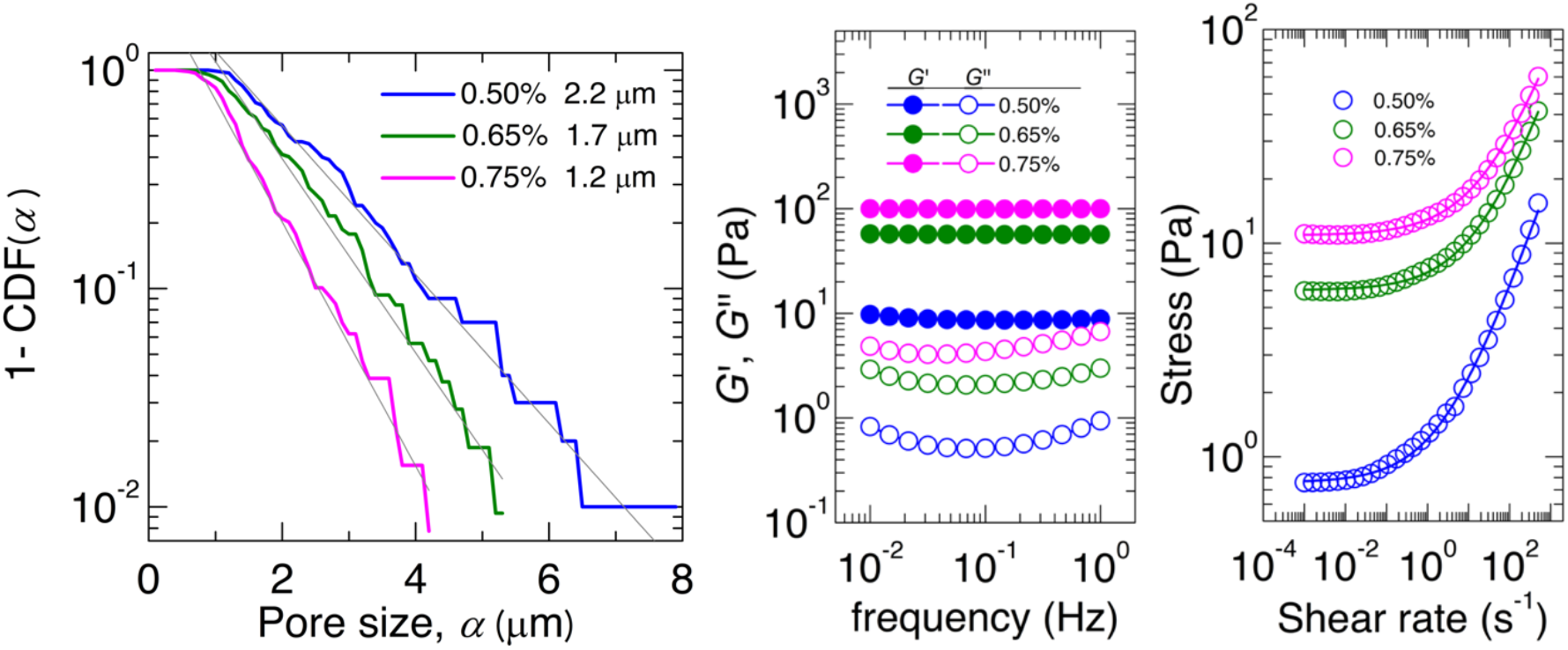
Characterization of porous media. (Left) Using measurements of many different tracers, which obtain the complementary cumulative distribution function 1-CDF(***α***) for each of the three porous media; here, 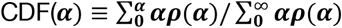 and *ρ*(***α***) is the number fraction of pores having dimension *α*. To determine the mean pore size, we fit an exponential ~***e^−α/A^*** to 1-CDF(***α***) for each medium and report the mean pore size as ***a = A + α***_0_ where ***α***_0_ is the largest pore size with CDF = 0. **(Middle)** Small-amplitude measurements of elastic (closed symbols) and viscous (open symbols) shear moduli, *G’* and *G”* respectively, of the porous media at different oscillatory frequencies. The legend indicates the hydrogel mass fraction used to prepare the media. The elastic moduli are frequency independent and are larger than the viscous moduli, indicating that the porous media are elastic solids. **(Right)** Measurements of shear stress as a function of applied unidirectional shear rate. At low shear rates the porous media behave like elastic solids with a yield stress given by the low shear rate asymptote of the curves shown; at high shear rates the media become fluidized.

**Figure S2:**
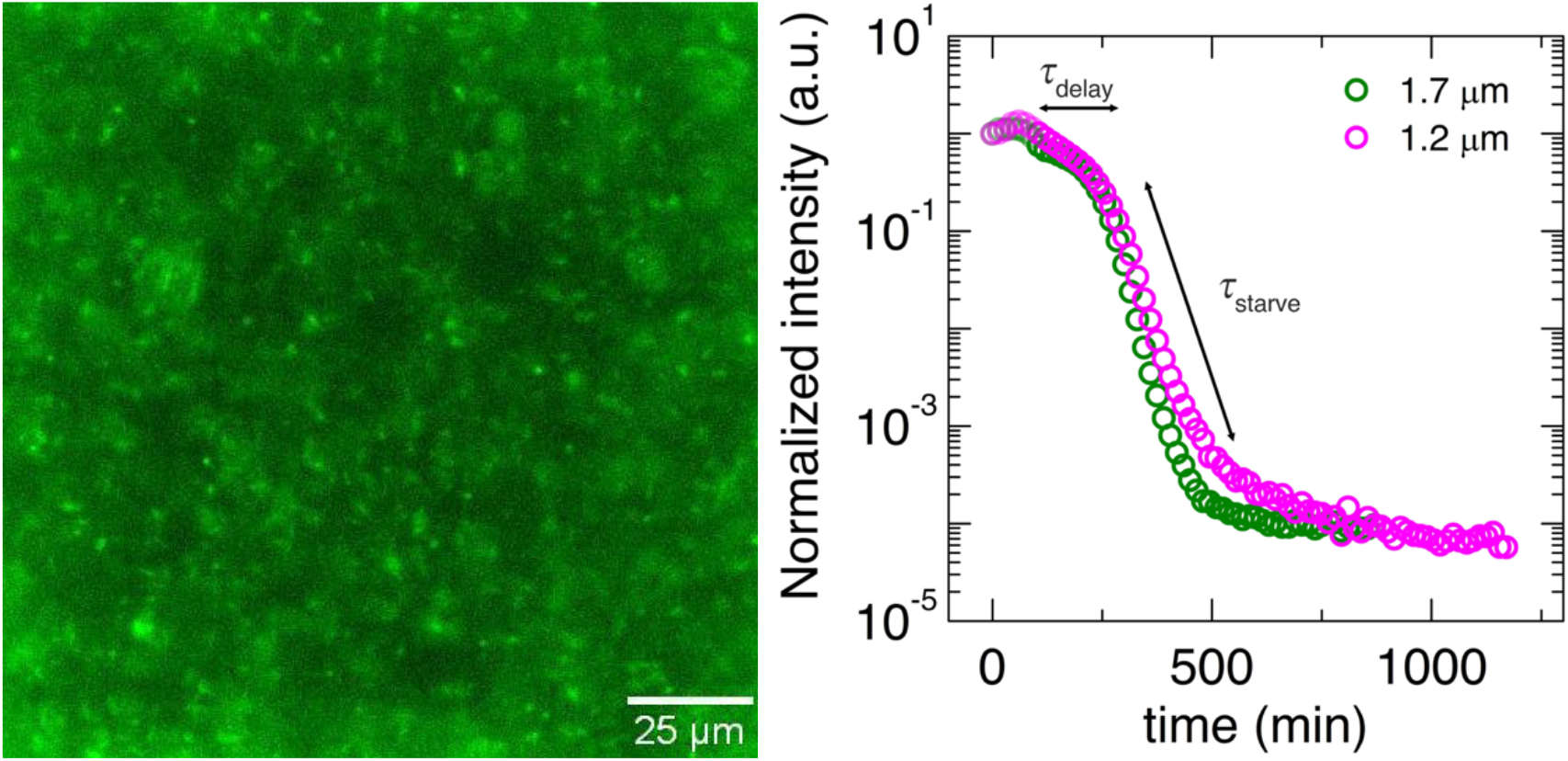
Nutrient-depleted cells remain trapped in initial position and eventually lose fluorescence. **(Left)** Fluorescence micrograph of cells trapped at the initial position of a cylindrical population ~17 h after 3D printing, well after front formation and propagation away. The cells are nonmotile and lose fluorescence; this image was taken by increasing the excitation intensity by ~29 times than used in all other experiments. **(Right)** Total fluorescence intensity measured at the center of a cylindrical population, normalized by initial intensity, in two different porous media with different mean pore sizes, indicated by the legend. Initially the cells migrate radially outward, leading to an initial slight decrease in fluorescent intensity (light symbols). The cells that remain trapped in their initial position maintain fluorescence for ***τ**_delay_*= 2 h and then lose fluorescence over a time scale of ***τ**_starve_*= 29.7 min.

**Figure S3:**
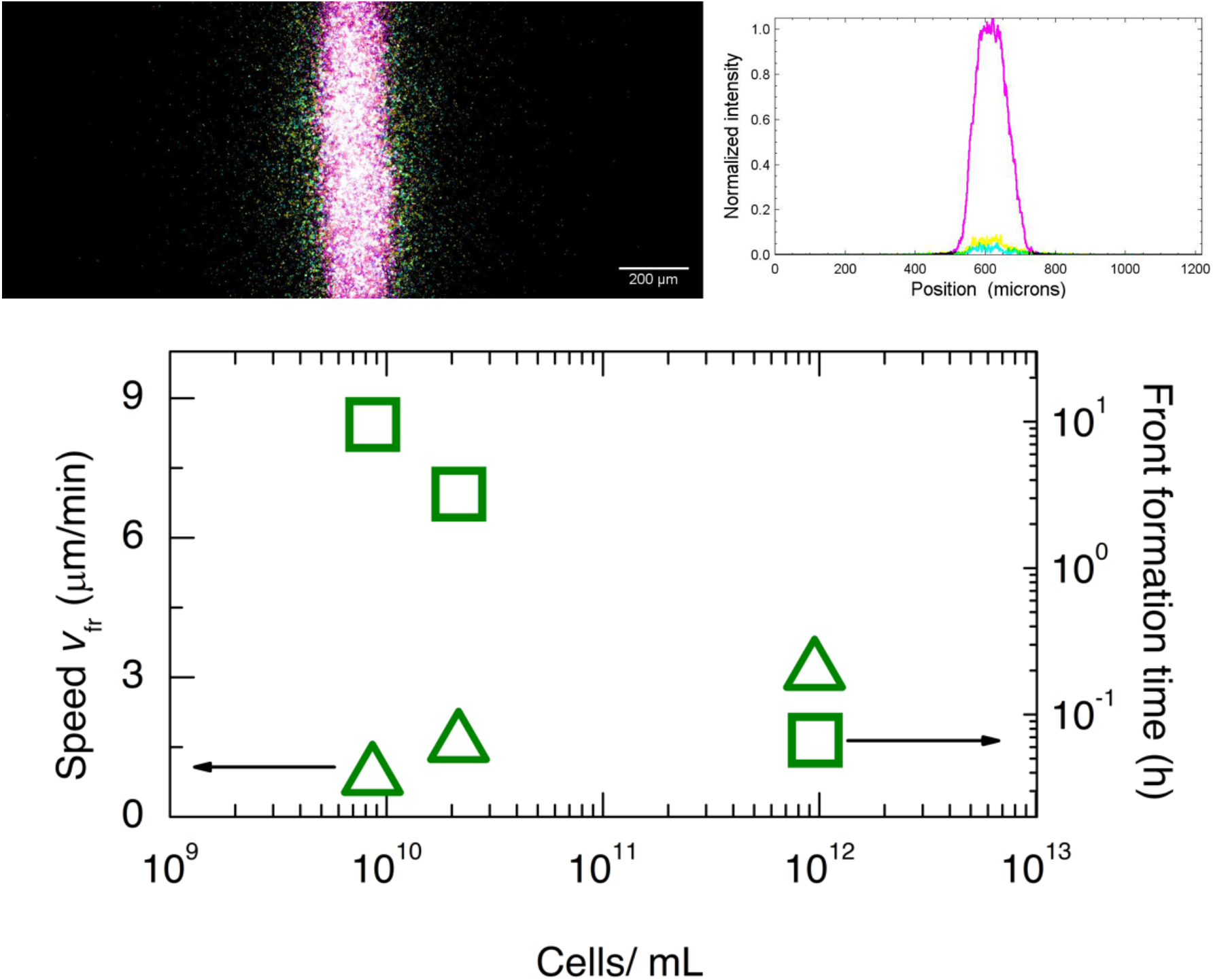
Front propagation is modulated by nutrient consumption. **(Top row)** Fronts do not form without nutrient. We 3D-print a cylinder of dense-packed *E. coli* within a medium prepared using 0.75% mass fraction hydrogel swollen in nutrient-free M9 minimal media instead of nutrient-rich EZ Rich media. We find that the population spreads outward diffusively, but does not form fronts, over the time scale required for fronts to form in nutrient-rich media (magenta: 0 h, yellow: 1 h, cyan: 2 h). Left panel shows bottom-up fluorescence micrograph of the population as it spreads, while right panel shows normalized intensity profiles through the 3D-printed population. **(Bottom row)** Front formation takes longer in more dilute populations. Data for a cylindrical population 3D-printed at different initial cell densities (horizontal axis) in a medium with *a* = 1.7 μm. The left vertical axis indicates the long-time speed of the propagating front formed ***v_fr_***, while the right vertical axis indicates the front formation time measured by monitoring when the fluorescence intensity measured from cells within the initial position of the cylinder begins to drop.

**Figure S4:**
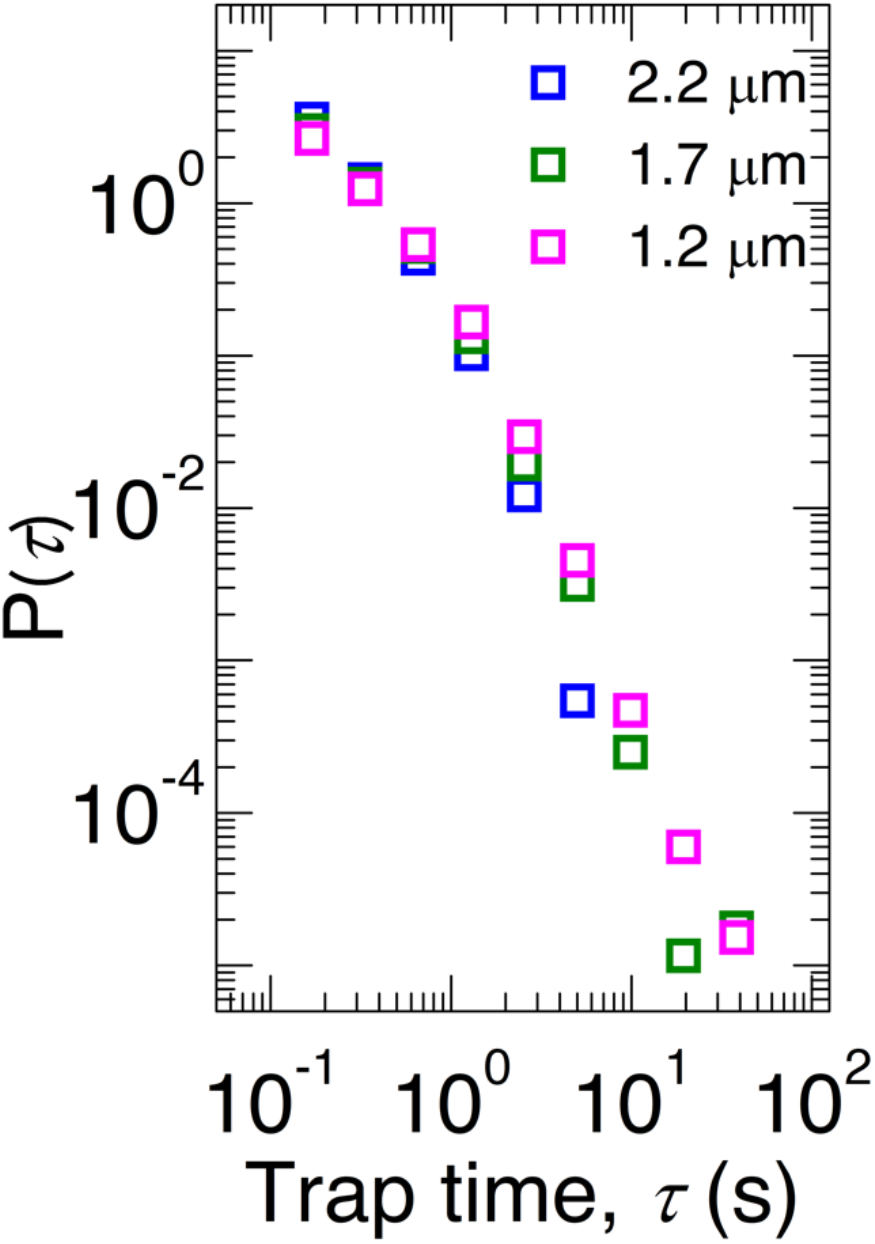
Trapping duration distributions of cells at the leading edge of the front. We track individual cells at the leading edge of propagating fronts measured in media with three different mean pore sizes, indicated by the legend. The cells exhibit hopping-and-trapping motility; similar to isolated cells in gradient-free media, the probability density of trapping durations exhibits a power-law tail. Notably, the majority of trapping durations is smaller than the mean chemotactic memory time of *E. coli,* ~4 s.

**Figure S5:**
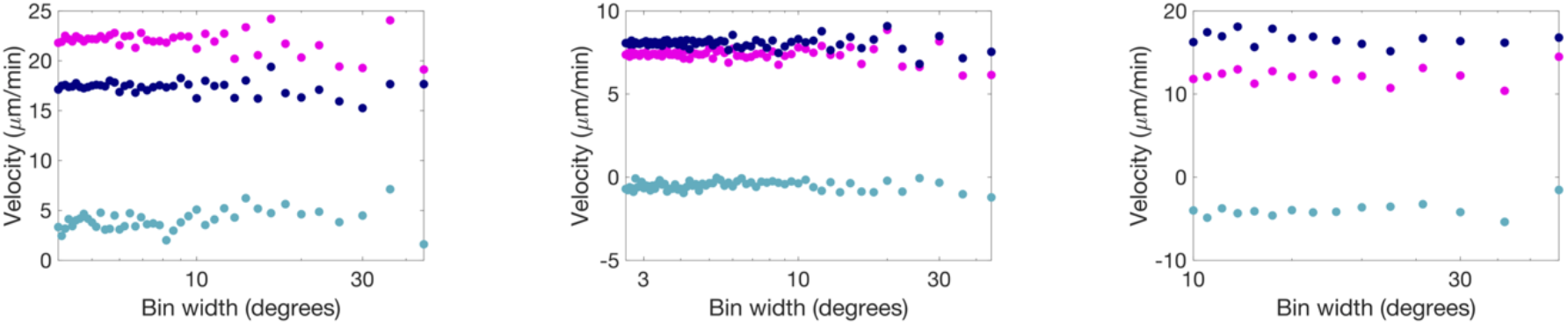
Bias in hopping orientation is the primary contributor to chemotactic migration. Plots show migration velocity calculated using the discrete sum version Eq. 1 of the main text, as described in the *Methods* text section “connecting single-cell motility to front propagation”, incorporating all factors (magenta), replacing orientation-dependent hopping lengths with the mean (dark blue) or replacing orientation-dependent hopping probability with a uniform distribution (teal). From left to right, data for the largest, intermediate, and smallest pore sizes are shown for different choices of the bin width, showing that the result reported in Fig. 3 is not sensitive to the choice of bin width. The data show that removing the bias in hopping orientation makes the largest difference in the calculated migration velocity for all pore sizes—that is, the bias in hopping orientation is the primary contributor to chemotactic migration.

**Figure S6:**
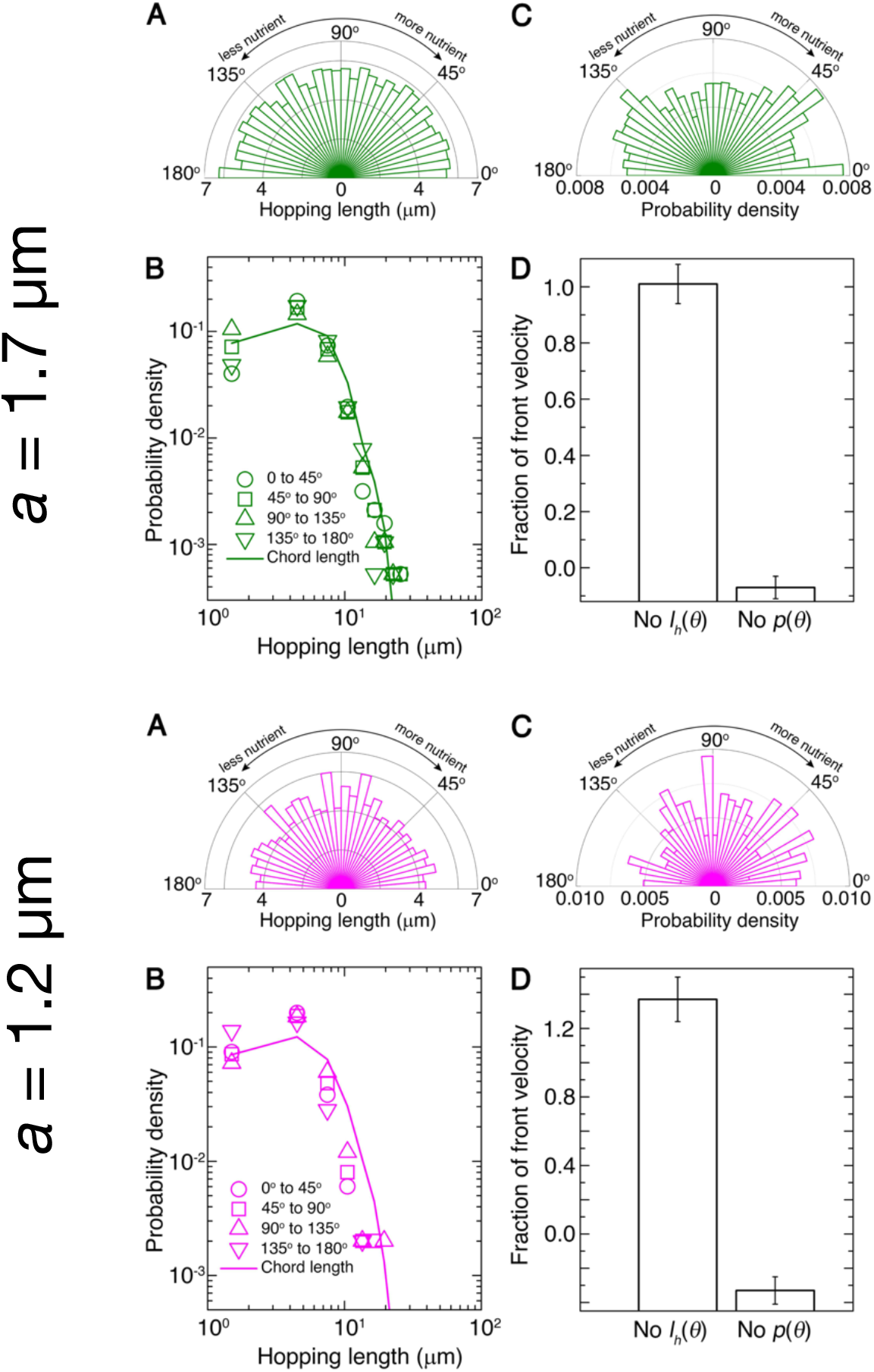
Biased motion of single cells in medium- and small-pore media. **(A)** Mean lengths of hops along different orientations |***θ***| with respect to the propagation direction. **(B)** Symbols show probability density of hopping lengths along different orientations within the ranges indicated by the legend; curve shows measured chord length distribution function for the porous medium. **(C)** Probability density of hopping along different orientations. **(D)** Chemotactic migration velocity calculated using Eq. 1, replacing orientation-dependent hopping lengths with the mean (first bar) or replacing orientation-dependent hopping probability with a uniform distribution (second bar). The second bar is slightly negative, indicating that measured hop lengths are on average slightly larger opposite the propagation direction–likely due to limited statistics. That front propagation would halt entirely or reverse without a bias in hopping orientation demonstrates that this bias is the primary driver of chemotactic migration. Error bars show standard deviation of velocity calculated using different angle bin widths.

**Figure S7:**
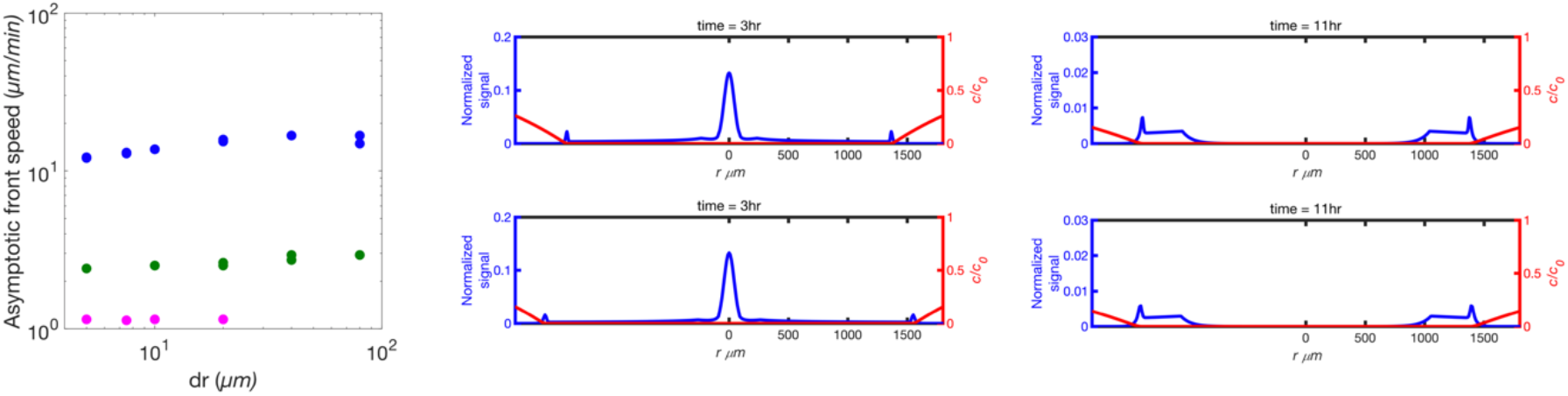
Choice of discretization in numerical simulations does not influence results. **(Left panel)** Long-time front speed for simulations representing chemotactic migration in porous media with *a* = 2.2, 1.7, and 1.2 μm (blue, green, magenta, respectively). We use two values of temporal discretization *dt* for each value of spatial discretization *dr* to ensure a sufficient time resolution was chosen; top and bottom rows show *dr*= 5 and 10 μm, respectively. Simulations showing normalized cell signal (blue) and normalized nutrient concentration (red) for *a* = 2.2 and 1.7 μm (**middle** and **right**panels, respectively) show minimal variation between choices of discretization indicated.

**Figure S8:**
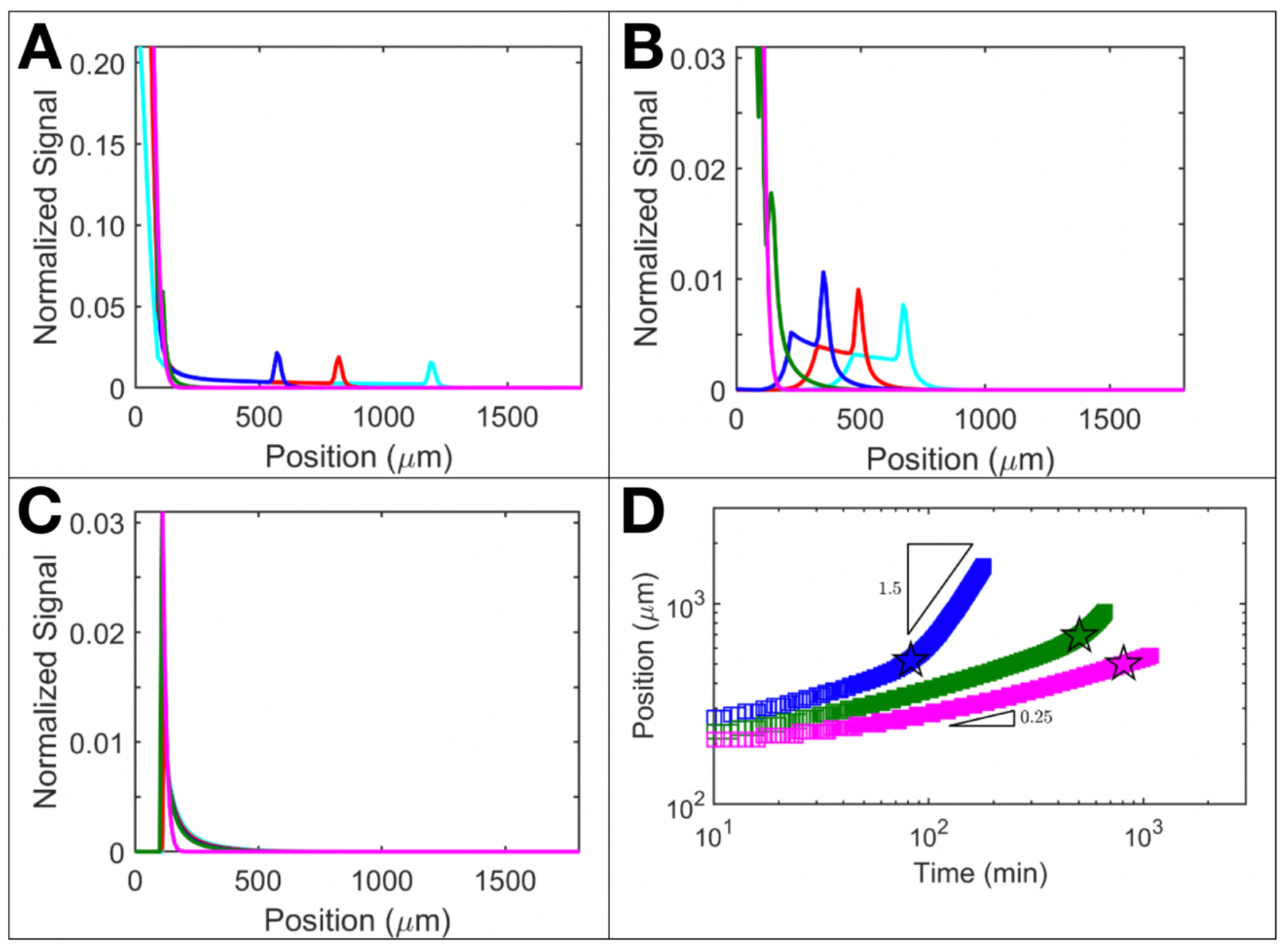
In the absence of growth, fronts still form and propagate, but are hindered in a confinement-dependent manner. Additional simulations of chemotactic migration without bacterial growth; panels **A-C** correspond to the same simulations as in Fig. 4A-C, but with; =0; colors correspond to the same times as in Fig. 4A-C. In the media with largest pores (A), front propagation appears to be similar to the case of non-zero growth, indicating that chemotaxis plays a dominant role in driving front propagation in these media; compare panel A to Fig. 4A. In the media with intermediate sized pores (B), front propagation is slower without growth; compare panel B to Fig. 4B. In the media with smallest pores (C), propagating fronts do not appreciably form over the simulation time scale, indicating that growth plays a dominant role in driving front propagation in these media; compare panel C to Fig. 4C. The resultant dynamics of the position of the leading edge of the front are shown in **(D)**.

**Figure S9:**
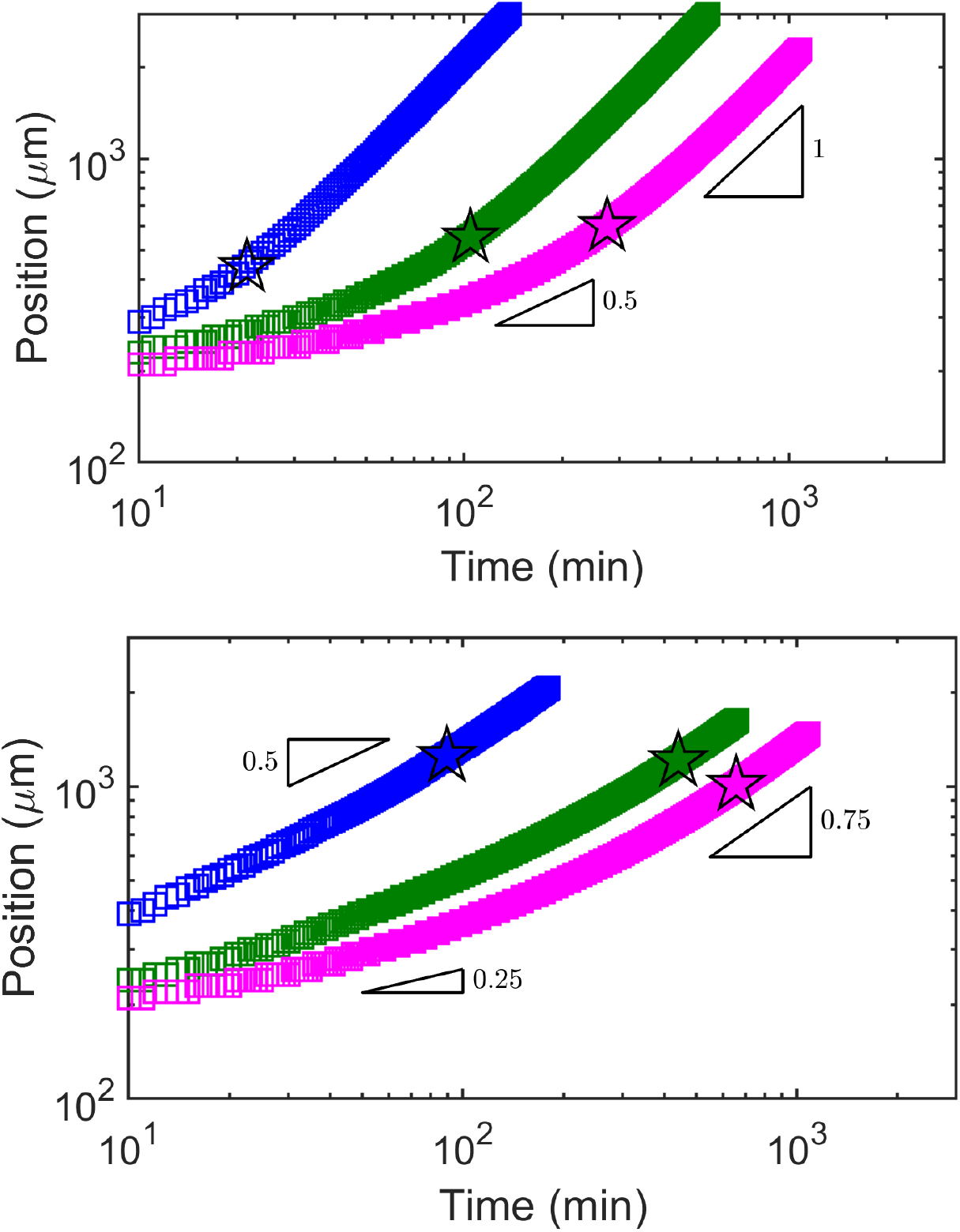
Further simulations of front propagation. **(Top)** Front propagation in rectilinear coordinates. Leading-edge position of the propagating front over time for simulations performed in rectilinear coordinates; colors correspond to those in Fig. 4. Stars indicate the crossover from diffusive to ballistic motion. In this case, we observe closer agreement to the crossover from diffusive to ballistic observed in the experiments than simulations performed in cylindrical coordinates. We conjecture that this agreement reflects the influence of boundaries in the experiment: while the experiments initially have cylindrical symmetry, with the initial 3D-printed cylinder placed far from all boundaries, as fronts propagate, they begin to approach the bottom boundary of the imaging chamber. Specifically, the simulations indicate that the region of nutrient depletion reaches the bottom boundary after ~0.5-1 h; in this case, the symmetry of the fronts is no longer cylindrical in the experiments, but has a rectilinear component. **(Bottom)**Without collisions, simulations do not exhibit the transition to ballistic motion observed in the experiments. To assess the importance of cell-cell collisions in the model, we perform the same simulations as in Fig. 4, but without the corrections to the motility parameters that incorporate cell-cell collisions. We fit the chemotactic parameter such that the speed over the last 30 minutes of the simulation matches the experiment, similar to our method for the main text, except here we impose no effects of cell-cell collisions. The values of chemotactic parameter obtained are 10, 0.9, and 0 μm^2^/s for the pore sizes in decreasing order, notably smaller than the values obtained by considering collisions. Moreover, none of the simulations achieve ballistic scaling in the absence of collisions.

**Figure S10:**
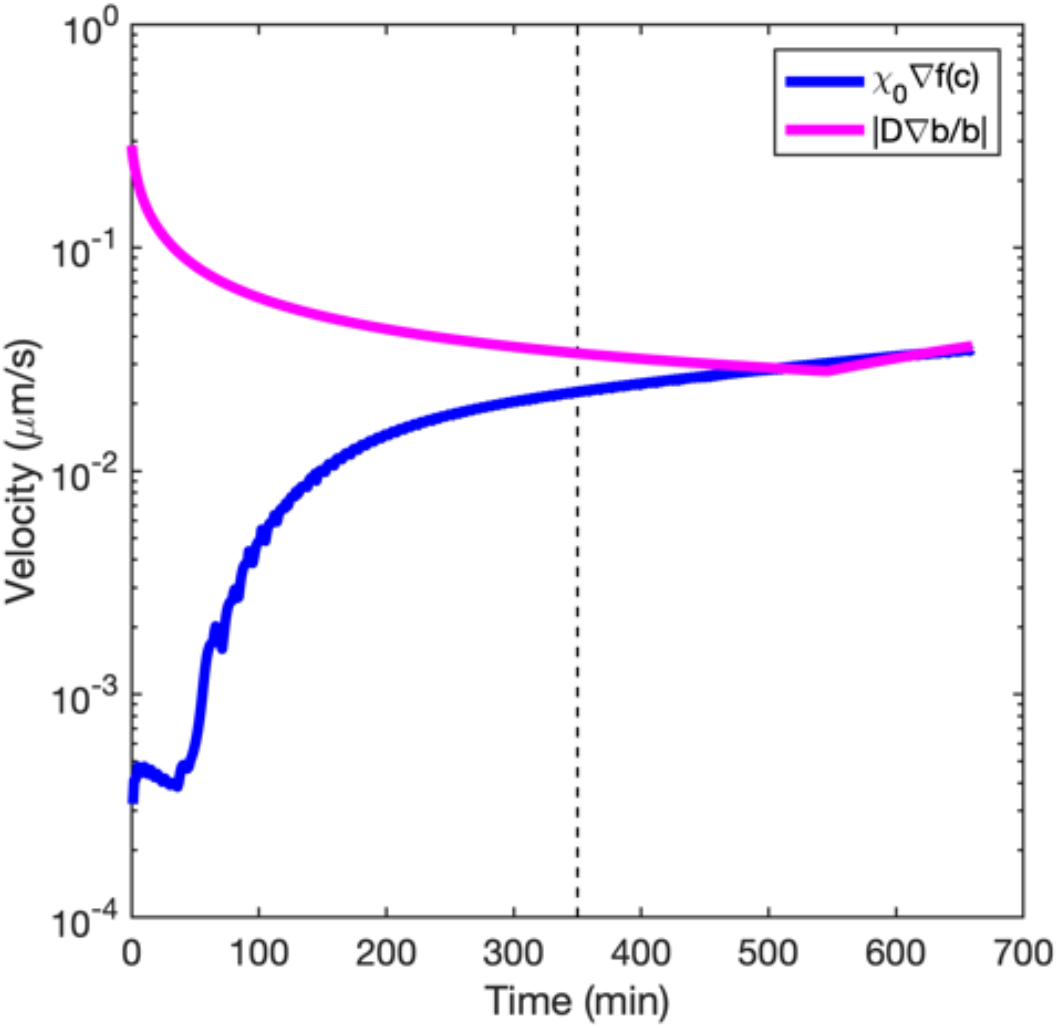
Simulations shed light on the physics underlying the observed diffusive-ballistic transition. An example for the case of intermediate pore-size media shows the variation of the maximum local chemotactic velocity (blue) and maximum local diffusive velocity (magenta) over time. At early times, the cell gradient is steep due to the sharp boundary of cells in the initial geometry of the population. This steep cell gradient drives diffusive flux, but decreases with time, as shown by the magenta curve. Meanwhile, chemotaxis begins low because (i) consumption must first reduce nutrient to within sensing levels and (ii) collisions halt the chemotactic response of cells within the dense starting region. Then, as the population spreads out, the chemotactic flux increases, and at the induction time (dashed line), the two velocities become comparable and eventually reach a steady state.

